# Global population genomics of two subspecies of *Cryptosporidium hominis* during 500 years of evolution

**DOI:** 10.1101/2021.09.09.459610

**Authors:** Swapnil Tichkule, Simone M. Cacciò, Guy Robinson, Rachel M. Chalmers, Ivo Mueller, Samantha J. Emery-Corbin, Daniel Eibach, Kevin M. Tyler, Cock van Oosterhout, Aaron R. Jex

**Author notes:** **Corresponding authors:** (ST), (CVO), (ARJ), (SMC). **E-mail addresses:** Swapnil Tichkule Simone M. Cacciò Guy Robinson Rachel M. Chalmers Ivo Mueller Samantha J. Emery-Corbin Daniel Eibach Kevin Tyler Cock van Oosterhout Aaron Jex.

## Abstract

Cryptosporidiosis is a major global health problem and a primary cause of diarrhoea, particularly in young children in low- and middle-income countries (LMICs). The zoonotic *Cryptosporidium parvum* and anthroponotic *C. hominis* cause most human infections. Here, we present a comprehensive whole-genome study of *C. hominis*, comprising 114 isolates from 16 countries within five continents. We detect two lineages with distinct biology and demography, which diverged circa 500 years ago. We consider these lineages two subspecies and propose the names *C. hominis hominis* and *C. hominis aquapotentis* (*gp60* subtype IbA10G2). In our study, *C. h. hominis* is almost exclusively represented by isolates from LMICs in Africa and Asia and appears to have undergone recent population contraction. In contrast, *C. h. aquapotentis* was found in high-income countries, mainly in Europe, North America and Oceania, and appears to be expanding. Notably, *C. h. aquapotentis* is associated with high rates of direct human-to-human transmission, which may explain its success in countries with well-developed environmental sanitation infrastructure. Intriguingly, we detected genomic regions of introgression following secondary contact between the subspecies. This resulted in high diversity and divergence in genomic islands of putative virulence genes (GIPVs), including *muc5* (CHUDEA2_430) and a hypothetical protein (CHUDEA6_5270). This diversity is maintained by balancing selection, suggesting a coevolutionary arms race with the host. Lastly, we find that recent gene flow from *C. h. aquapotentis* to *C. h. hominis*, likely associated with increased human migration, may be driving evolution of more virulent *C. hominis* variants.

## Main

Cryptosporidiosis is a leading cause of diarrhoea in children under five globally^1,2^, resulting in an estimated 48,000 deaths annually. Among parasitic diseases, it is second only to malaria in global health burden, with an overall impact of ∼12.8 million Disability Adjusted Life-Years (DALYs)^1^. Human cryptosporidiosis is primarily caused by *Cryptosporidium parvum*, a zoonoses common in young ruminants^3,4^, and *C. hominis*, which is anthroponotic and the more prevalent species globally^5^. The disease burden is overwhelmingly skewed to low and middle-income countries (LMICs)^6^, particularly in sub-Saharan Africa^1^, where *C. hominis* predominates. However, *Cryptosporidium* remains a significant public health problem in wealthy countries through large water or foodborne outbreaks and direct transmission in day-care facilities, hospitals and other institutions^7,8^. Presently, there are no vaccines or effective drugs to treat the infection. Hence, control depends on prevention of infection, which is driven by a strong understanding of the parasite’s epidemiology.

The global epidemiology of cryptosporidiosis varies by geographic region, socioeconomic status and a range of risk factors^6,9^. Understanding of this epidemiology is underpinned by molecular typing, based mainly on the highly polymorphic *gp60* gene^10^. This work has identified numerous genetic variants within each species and indicated a complex population genetic structuring. In *C. parvum*, population structure varies globally from clonal to epidemic to panmictic, likely due to varying ecologic factors^11-13^. Genetic variants found exclusively in humans point to an anthroponotic *C. parvum* lineage (IIc)^14,15^, with a recent genomic study recognising two subspecies, the zoonotic *C. parvum parvum* and the anthroponotic *C. parvum anthroponosum*^16^. Interestingly, this study found that both subspecies occasionally still hybridise and exchange genetic variation. These exchanges overlapped with similar genomic regions undergoing genetic introgression between *C. hominis* and *C. parvum anthroponosum*, indicating candidate sites underpinning adaptation to human-specific infection^16^. *Cryptosporidium hominis* appears to have a largely clonal population structure, dominated by specific variants in different regions^17^. The IbA10G2 *gp60* subtype, although found in LMICs^18^, is the dominant variant in high-income countries and accounts for up to ∼45% of all *gp60*-typed human infections^18^. This subtype is linked to most major waterborne outbreaks in wealthy countries for which genetic typing is available^19- 23^. The IbA10G2 subtype also appears more readily capable of direct human-to-human transmission^24^ and may cause more severe disease^25^.

Studies of *C. hominis* molecular epidemiology pose several essential questions that have major implications for global control of cryptosporidiosis. Specifically: (1) what is the IbA10G2 subtype, and is this *gp60*-defined subtype reflective of a phylogenomically divergent *C. hominis* lineage that predominates in wealthy countries; (2) if so, does this lineage undertake reduced levels of genetic recombination with global *C. hominis* populations; (3) and can signatures within the genomes IbA10G2 typed isolates identify its taxonomic status, reveal the factors underpinning its increased virulence and its dominance in wealthy countries, and identify its influence on global parasite population structure? To address these questions, we performed a global study of 114 *C. hominis* genomes, comparing the IbA10G2 subtype to published genomes representing locally acquired infections from 16 countries across five continents. The insights gained from these analyses are particularly relevant for public health; the IbA10G2 subtype has already been identified as an emerging host-adapted, virulent, and hyper-transmissible population, making it a threat to human health in high-income countries^26-28^.

## Results

### Genomic evidence of population sub-structuring at the continental level

All 114 *C. hominis* isolates included in this study were confirmed as single variant infections (estimated MOI=1). We identified 5,618 biallelic SNPs among these samples and used these to explore the population structure of *C. hominis*. Our analyses identified two major clusters (Fig. 1a), separating European, North American and Oceanian samples from Asian and African samples. We also saw minor clustering separating African from Asian samples. STRUCTURE analysis provided further support for these findings (Fig. 1b). Overall, ∼94% of isolates clustered geographically. Exceptions included a small number of infections (e.g., UKH30 (from UK) and Sweden_SRR3098109 (from Sweden) acquired during international travel. STRUCTURE analysis also identified a fourth cluster, including three African and one European isolate, with unique population ancestry (cluster 4 in Fig. 1b).

**Fig 1.**
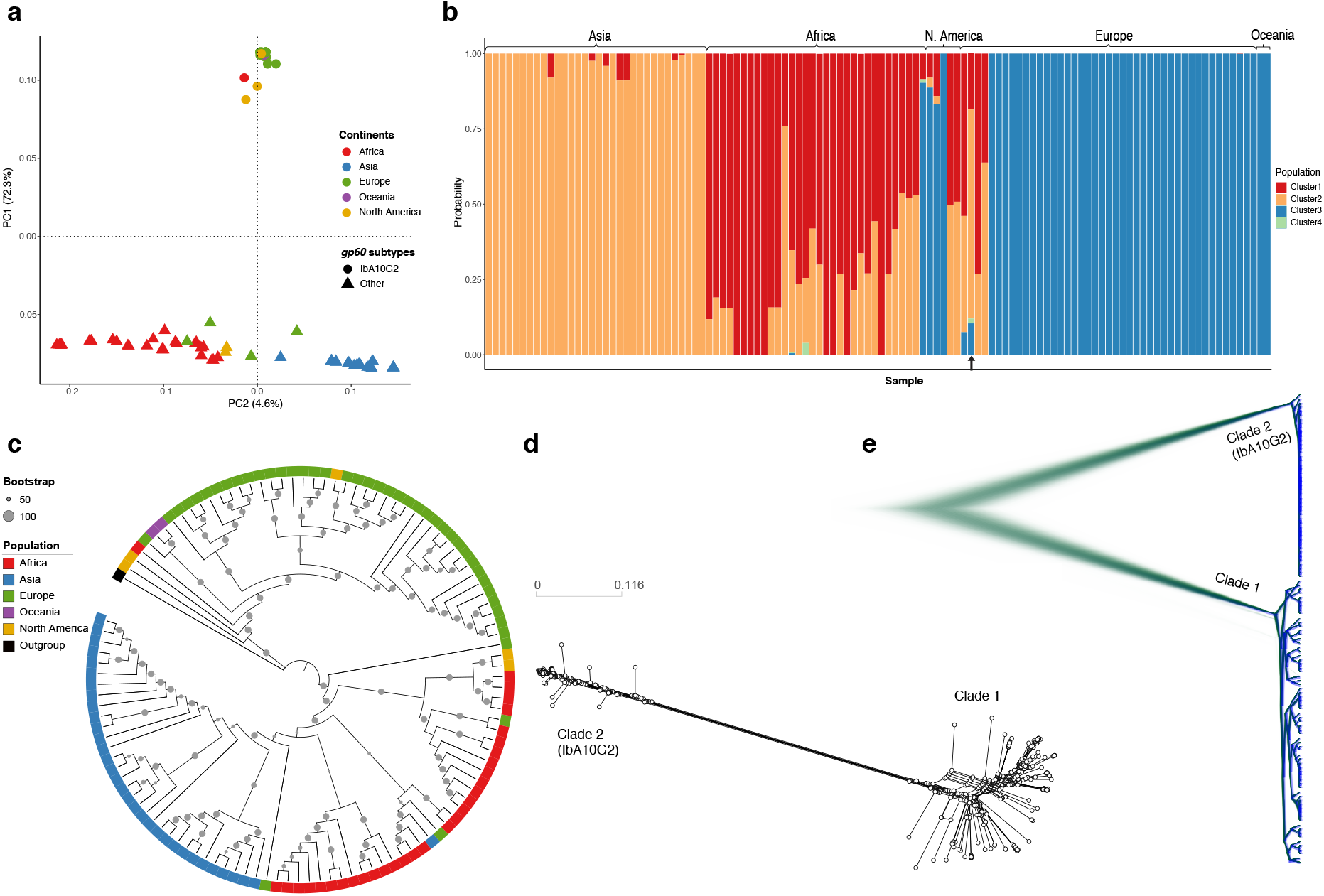
Global population structure of *C. hominis* isolates illustrating their sub-structing and diversification. **a**. PCA of isolates based on their whole genome SNPs, highlighting three clusters of isolates which are predominately based on continents of origin. Isolates were color coded with their continent of origin. Isolates associated with *gp60* subtype IbA10G2 were represented with solid circles while non-IbA10G2 with solid triangles. **b**. Structure plot illustrating population genetic ancestry and the admixed nature of the *C. hominis* isolates. The plot was obtained assuming an optimum value of K=4. The black arrow (bottom) indicates the highly admixed isolate (UK_UKH4), which includes all four ancestries. **c**. Phylogenetic clustering demonstrating two clades. **d**. Splitstree and **e**. Densitree are also demonstrating two major clades comprising isolates, which are separated based on low- and high-income countries of origin, respectively. *Cryptosporidium h. aquapotentis* includes isolates associated with *gp60* subtype IbA10G2 while *C. h. hominis* includes isolates associated with other *gp60* subtypes.

### Genomic evidence of population diversification

Maximum Likelihood (ML) analysis (Fig. 1c) identified two major clades, one corresponding to Asia and Africa (clade 1) and the other to Europe, North America and Oceania (clade 2). All isolates within clade 2 had *gp60* subtype IbA10G2, and no IbA10G2 isolates clustered with clade 1. The two clades were estimated to have diverged 488 (84 – 2,199; 95% HPD) years ago. This divergence is supported by a network (Fig. 1d) and SplitTree and DensiTree (Fig. 1e) analyses. Considering the stability of these two genomically distinct lineages and evidence below of their reproductive isolation, we propose their recognition as separate subspecies. We propose clade 1, which comprises infections observed in low-to middle-income countries, mostly from Africa and Asia, be recognised as *C. hominis hominis*, referring to the fact that this subspecies represents the majority population. We propose clade 2 (IbA10G2 subtype) includes isolates from high-income countries, namely Europe, North America and Oceania, be named *C. hominis aquapotentis* (strong water), as it predominates in countries with longstanding high sanitation and water quality indices.

### Demographic histories

To understand the demographic histories and estimate the change in the effective population size (*Ne*) through time, we constructed a Bayesian Skyline Plot (BSP)^29^ for *C. h. hominis* and *C. h. aquapotentis*. Parasite isolates from low-income countries (*C. h. hominis*) experienced a marked population contraction recently from *Ne*=∼5000 to *Ne*=∼100 (Fig. 2a), which is supported by a higher proportion of positive Tajima’s D values (Fig. 2b). In contrast, the isolates from high-income countries (*C. h. aquapotentis, gp60* subtype IbA10G2) had a stable effective population size (*Ne*=∼1000) and a higher proportion of negative Tajima’s D values (Fig. 2c). The distribution of Tajima’s D in *C. h. aquapotentis* is significantly smaller than zero (one-sided t-test: t = -28.883, df = 681, p-value < 2.2e-16), which is consistent with recent population expansion. Simulation-based projections of the evolution of the overall *C. hominis* population finds it is being shaped by ‘recent gene flow’, likely indicating recent secondary contact (Fig. 2c-d) between the two subspecies ∼165 (24 – 647; 5-95% CI) years ago.

**Fig 2.**
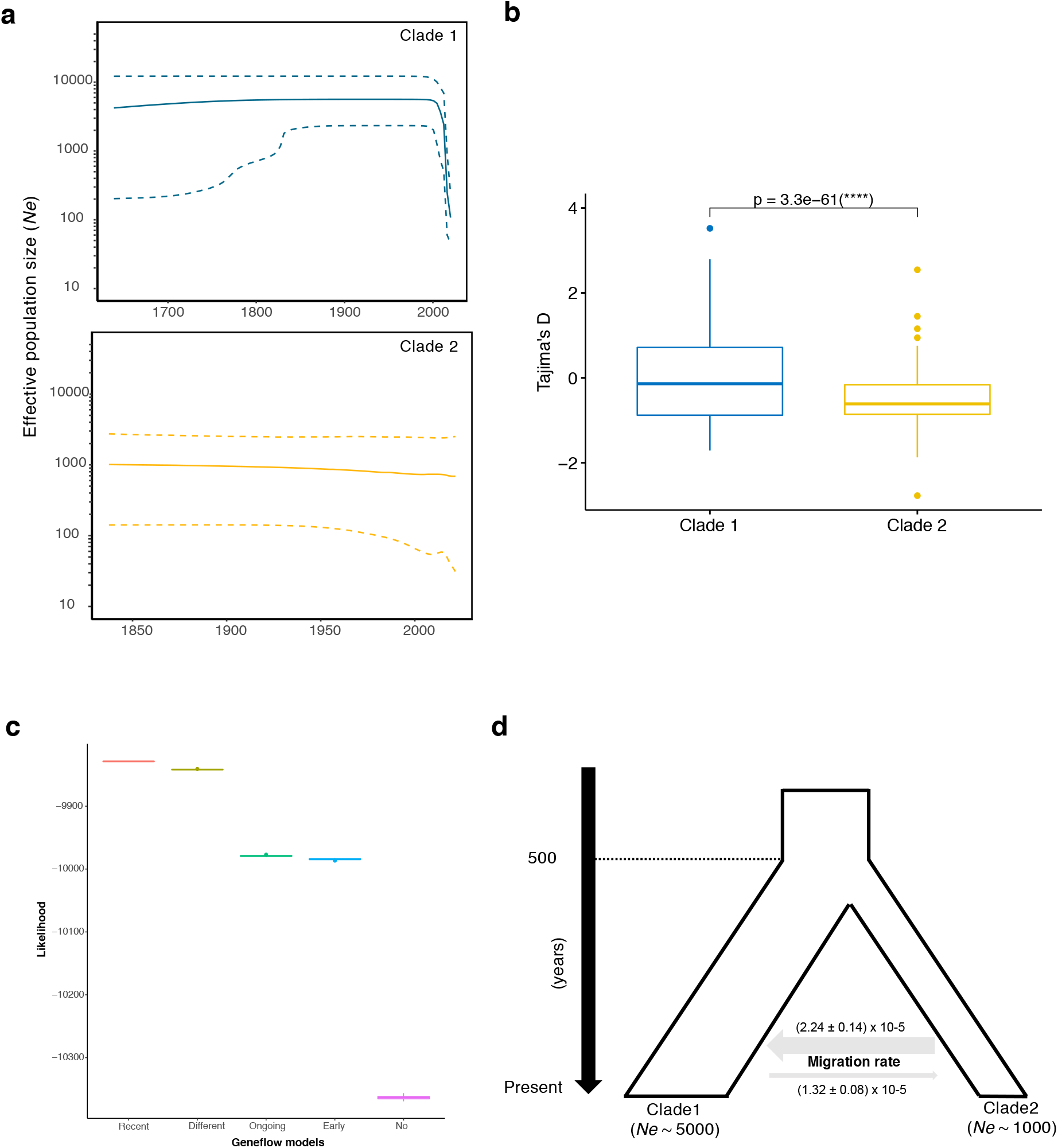
Demographic histories and population size and secondary contact between *C. h. hominis* (clade 1) and *C. h. aquapotentis* (clade 2). **a**. Bayesian Skyline plots (BSP) depicting change in *Ne* (effective population size) through time, for both the clades. The central dark line and the upper and lower dashed lines on Y-axis are mean estimates and 95% HPD intervals of *Ne*, respectively. X-axis is time in years, running backwards. **b**. Boxplot showing significant difference (two-sided t-test) in Tajima’s D values between *C. h. hominis* (clade 1) and *C. h. aquapotentis* (clade 2). **c**. Higher likelihood for “recent geneflow” model (in red). Comparing likelihood distributions of geneflow models and observed significant difference (one-way ANOVA test, F = 2629761, df = 4, p-value <2e-16). Further, Post-hoc Tukey-HSD test revealed difference in likelihood between all the models (p-value < 1e-16). **d**. Graphical representation of demographic history of *C. hominis*, illustrating recent secondary contact and migration rates between the two clades (mean ± SE).

We found evidence of two selective sweeps on chromosome 6 in *C. h. hominis* based on composite likelihood ratio (CLR) statistic using SweeD^30^ (Supplementary Fig. 1). However, we do not see any other hallmarks of a selective sweep in these regions based on π or Tajima’s D (see Supplementary Fig. 2). Possibly, the selective sweeps were incomplete or occurred in the distant past, eroding their genomic signatures. Nevertheless, this may have contributed to the recent decline in effective population size of *C. h. hominis* (Fig. 2a); genetic variation could have been lost during the selective sweep not only in the affected chromosomal regions, but throughout the genome in this largely clonally reproducing organisms, as the selectively favoured variant replaced other existing variants.

### Linkage, recombination, introgression and gene flow between subspecies

#### Distinct patterns of decay of linkage disequilibria

We performed independent linkage analyses for each subspecies to infer the recombination rate within parasite populations from low- and high-income countries (Fig. 3a). *Cryptosporidium h. hominis* had more rapid linkage disequilibrium (LD) decay than *C. h. aquapotentis* (IbA10G2), consistent with genetic exchanges through gene flow and recombination. In contrast, the strong LD in *C. h. aquapotentis* (IbA10G2) supports our hypothesis of a recent population expansion (see below).

**Fig 3.**
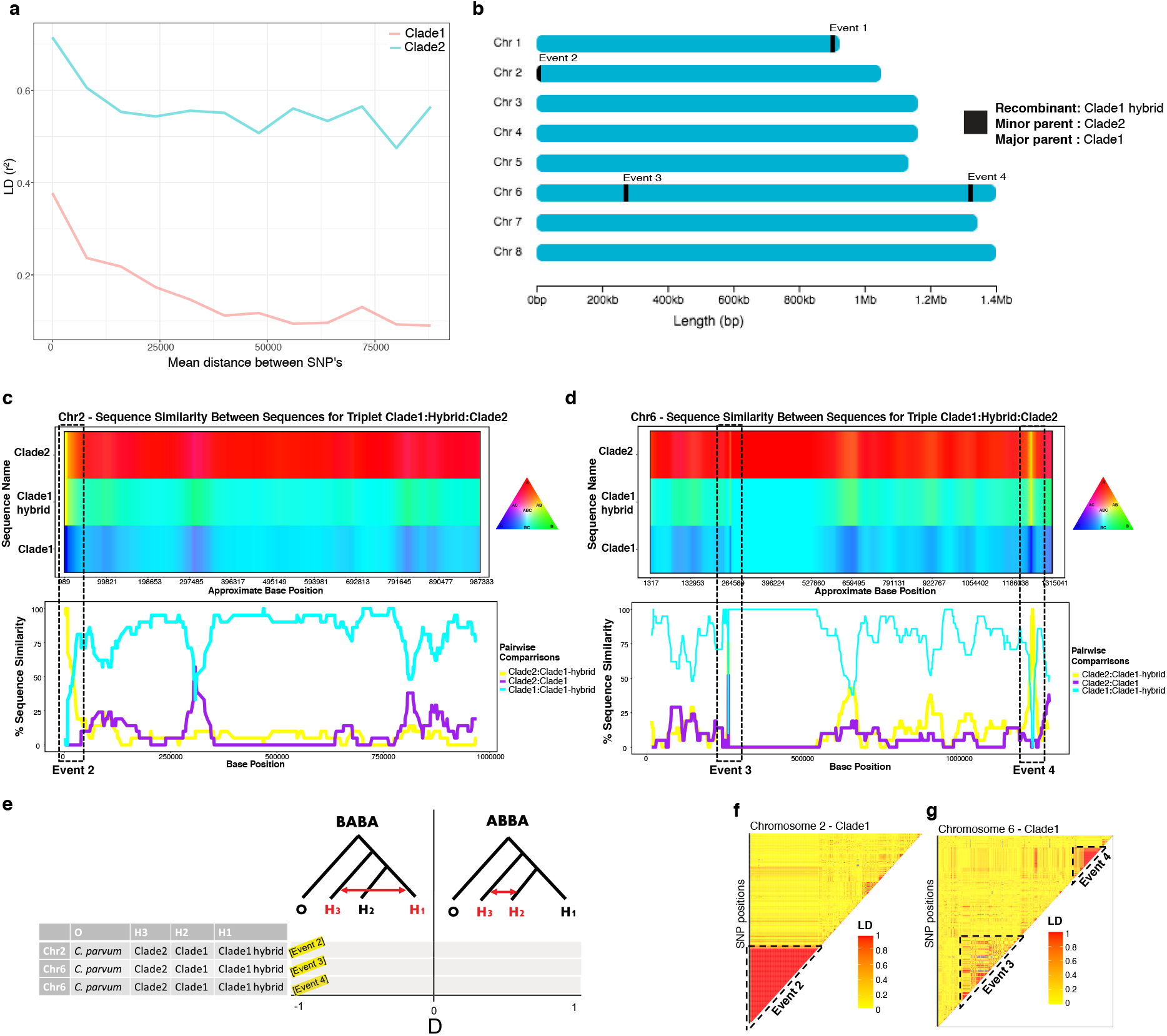
Analyses of recombination and gene flow between *C. h. hominis* (clade 1) and *C. h. aquapotentis* (clade 2). **a**. Linkage disequilibrium (LD) decay plot showing rapid decay of linkage between SNPs in *C. h. hominis* (clade 1) compared to *C. h. aquapotentis* (clade 2). **b**. Graphical representation of recombinant breakpoint positions detected by RDP4 program between *C. h. hominis* (clade 1) and *C. h. aquapotentis* (clade 2). **c-d**. HybridCheck plots representing genomic signature of introgression in chromosome 2 and 6, respectively. Analysis for chromosome 1 was excluded due to unknown parental sequences. The plots were generated for random set of triplets that includes recombinant (hybrid), minor (donor) and major (recipient) parental sequence, as detected by RDP4 program. Introgressed blocks (recombinant breakpoints) were illustrated with dashed boxes, showing high similarity between the recombinant (*C. h. hominis* hybrid isolates) and minor parent (*C. h. aquapotentis* isolates). **e**. Gene flow analyses with ABBA-BABA test, representing D statistics for the random sets of triplets (as used in c-d) along with *C. parvum* as an outgroup. D statistic values close to -1 at all three recombinant events, suggesting geneflow between H1 and H3. **f-g**. Pairwise LD of SNPs in chromosomes 2 and 6 of *C. h. hominis* showing red-blocks of high linkage between SNPs in introgressed events 2-4.

#### Recombination and regions of secondary contact

Using RDP4^31^, we identified significant recombination events between the two clades in chromosome 1 (event 1), 2 (event 2) and 6 (event 3 and 4) (Fig; 3b and Supplementary Table 2). This indicated secondary contacts between *C. h. hominis* and *C. h. aquapotentis* (IbA10G2), which resulted in rare genetic exchanges between these otherwise diverged clades. We also undertook additional analyses of the highly admixed European isolate (UK_UKH4 of *gp60* subtype IaA14R3) that showed unique ancestry (Fig 1B) and clustered with low-income countries (*C. h. hominis*) (see Supplementary Text).

#### Signature of introgression and gene flow between the clades

We analysed the recombination events in more detail to better understand the implications of genetic introgression between the two subspecies. Determining the signature of genetic introgression is crucial as these regions can also be responsible for increasing genetic diversity and providing novel substrate for natural selection in host-parasite coevolution^32^. We used HybridCheck^33^ to perform introgression analyses for recombinant events 2 – 4 (event 1 was excluded due to a missing parental sequence). We randomly selected a triplet (recombinant, minor parent and major parental lines) from the RDP output (Supplementary Table 2), which revealed a clear signature of introgression (Fig. 3 c-d), also supported by an ABBA-BABA test (Fig. 3e). Additionally, we calculated the pairwise r^2^ between SNPs within chromosomes of *C. h. hominis* to assess linkage among SNPs in the introgressed regions (Fig. 3f-g). Large blocks of high LD that encompass the introgressed regions suggested each had been exchanged as a single event between *C. h. hominis* and *C. h. aquapotentis* (IbA10G2) ∼165 (24 – 647; 5-95% CI) years ago.

### Could recent introgression increase virulence?

Our analyses suggest *C. h. hominis* and *C. h. aquapotentis* have been diverged and largely reproductively isolated for ∼500 years. Noting that the latter subspecies has come to dominate infections within high-income countries^34^, is more virulent^25^, appears better able to transmit through direct human-to-human contact^24^ and possibly at a lower infectious dose^22^, *C. h. aquapotentis* may owe its success to being better adapted to human infection. If this is the case, it is possible that recent introgression between these two subspecies could select for more virulent and transmissible *C. hominis* subspecies in low-income countries, particularly noting the recent genetic bottlenecking we have observed in *C. h. hominis* within the last decades.

#### Identification of potential virulence genes

To explore this, we first predicted candidate virulence genes in *C. hominis* likely to be involved in the host-parasite interaction and engaged in a coevolutionary arms race with the host. Broadly, such genes are under continuous adaptation and involved in reciprocal, adaptive genetic changes while interacting with host^32^. During coevolution, evolutionary forces act on virulent genes to create genetic variation through mutation, recombination and gene flow and moulds genetic variation through selection. To identify putative virulence genes in *C. hominis*, we selected the top 5% most highly polymorphic genes based on nucleotide diversity (π). These were filtered for the top 25% of genes under balancing selection, based ranked Tajima’s D. Finally, these genes were filtered by selecting the top 50% of genes with the highest proportion of non-synonymous mutations, based on ranked Ka/Ks ratio. Using this approach, we identified 24 highly polymorphic genes (Supplementary Table 4) that are rapidly mutating at the protein level and under selective pressure. These genes were significantly more polymorphic (two-sided t-test: t = -2.9062, df = 23, p-value = 0.007957), under stronger balancing selection (two-sided t-test: t = -10.393, df = 23, p-value = 2.533e-10) and positive selection (two-sided t-test: t = -2.4736, df = 23, p-value = 0.021) compared to all remaining polymorphic genes. Moreover, these genes are enriched in recombination regions (χ2= 225.04, df = 1, p-value = 0.00049), showing that gene flow within *C. hominis* is inclined towards elevated levels of genetic variation. Overall, these genes are enriched for extracellular (χ2= 13.608, df = 1, p-value = 0.00349) and signal peptide proteins (χ2= 6.69, df = 1, p-value = 0.015), which is consistent with their potential relevance for host-parasite interaction. Together, these results provided the evidence of genomic islands of putative virulence genes (GIPVs: Fig. 4) that are likely to be in a coevolutionary arms race with the host, undergoing frequent recombination and accumulating beneficial polymorphisms that are under selection, and that this increased population diversity.

**Fig 4.**
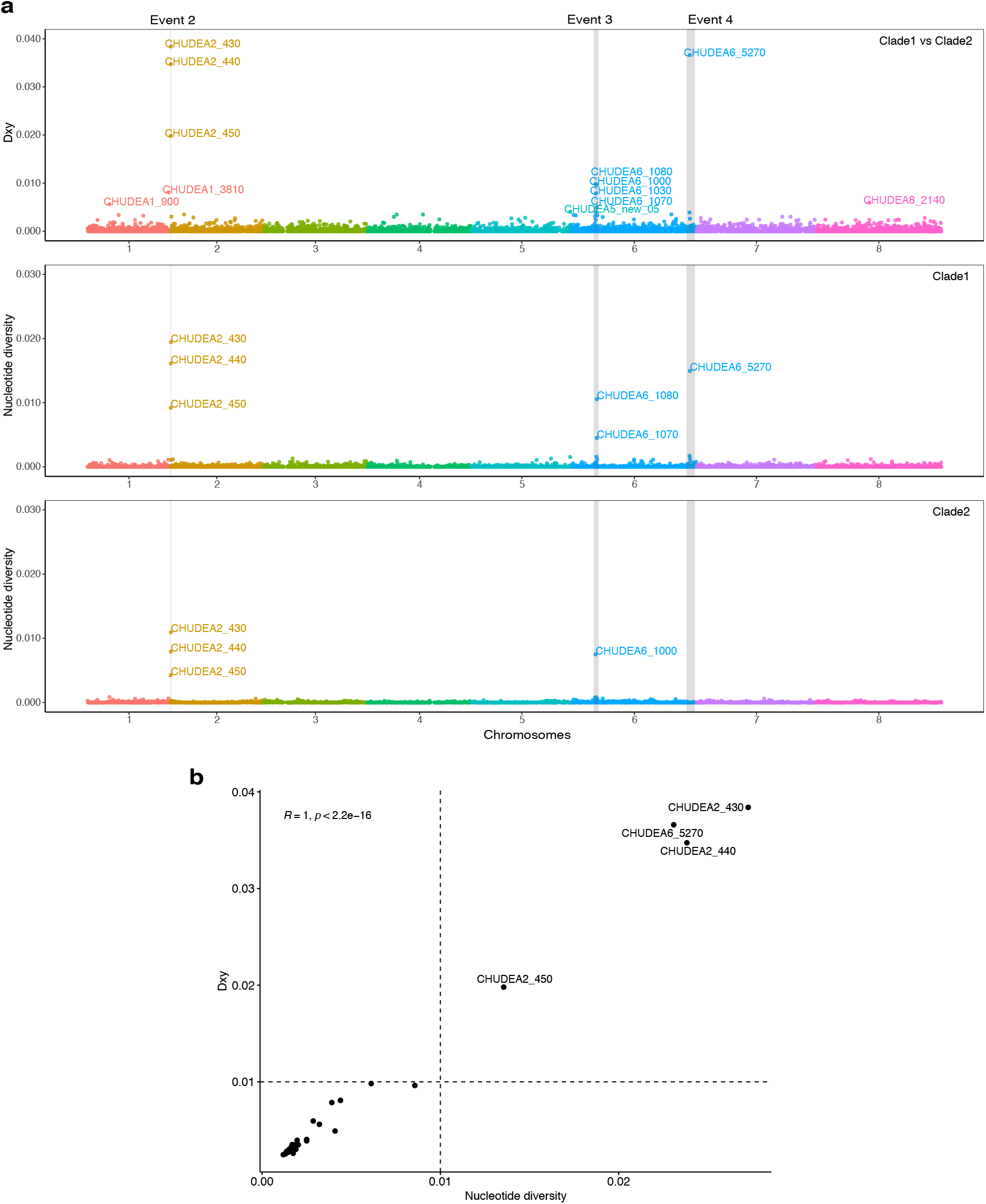
Population genetic analyses of genomic islands of putative virulence genes (GIPVs). **a**. Population genetic and divergence analyses of introgressed regions. X-axis represents genomic positions of eight chromosomes highlighted with different colours. Population divergence (Dxy) between *C. h. hominis* (clade 1) and *C. h. aquapotentis* (clade 2) for each gene were plotted on Y-axis (top panel). Nucleotide diversity (π) for *C. h. hominis* (middle panel) and *C. h. aquapotentis* (bottom panel) for each gene, were also plotted on Y-axis, respectively. The breakpoints of four recombination events (event 1-4) were indicated by grey vertical boxes. Event 1 was un-detected in *C. h. aquapotentis*. **b**. Correlation between π and Dxy were plotted to identify polymorphic and potential virulent genes.

#### Subspecific divergence of putative virulence genes

We investigated whether any virulence genes predicted above were highly diverged between *C. h. hominis* and *C. h. aquapotentis* and under diversifying selection between the two subspecies. To do this, we calculated absolute divergence (Dxy), representing the average proportion of differences between all pairs of sequences between *C. h. hominis* and *C. h. aquapotentis* (IbA10G2), revealing differential gene flow during their reproductive isolation (Fig. 4a). We then calculated the correlation between diversity (π) and divergence (Dxy) for each gene (Fig. 4b). This approach identified four clear outlier genes, which were the most divergent, diverse, and rapidly mutating at the protein level. These genes encoded three mucins (CHUDEA2_430, CHUDEA2_440 and CHUDEA2_450) arrayed in a cluster on chromosome 2, and a hypothetical protein (CHUDEA6_5270) found on chromosome 6 (Fig. 4b, Supplementary Table 4). The *gp60* gene ranked 5^th^ in the list of virulence genes but does not sit as a clear outlier from the other genes identified in our assessment. Two of these genes, CHUDEA2_430 (LRT, p-value = 0.005) and CHUDEA6_5270 (LRT, p-value = 0.017), are under statistically significant diversifying selection between *C. h. hominis* and *C*.

#### h aquapotentis

Lastly, we looked at each codon position within each of the four outlier genes to detect codon specific diversifying selection, which might be overlooked in the overall gene. We detected episodic diversifying selection (LRT p-value < 0.05) at codon positions 92, 111 and 138 of CHUDEA2_430 and 97 and 105 of CHUDEA6_5270 using the Mixed Effects Model of Evolution (MEME) method implemented in Datamonkey^35^. These sites represent putative codons harbouring genetic polymorphisms that experience periods of strong diversifying selection. This strongly suggests CHUDEA2_430 and CHUDEA6_5270 are likely candidate virulence factors participating in a coevolutionary arms race and contributing to the divergence of *C. h. hominis* and *C. h. aquapotentis*.

#### Recent introgression of putative virulence genes

Finally, we asked whether the recent recombination events in chromosomes 2 and 6 between *C. h. hominis* and *C. h. aquapotentis* might include genes associated with increased virulence, indicating increasing virulence of *C. hominis* globally within the past decades, possibly linked to increased human migration. These introgressed regions had significantly elevated levels of nucleotide diversity (two-sided t-test: t = -3.0026, df = 27, p-value = 0.0057), divergence (two-sided t-test = -3.0043, df = 27, p-value = 0.0057) and balancing selection (two-sided t-test: t = -3.0125, df = 27, p-value = 0.0055). Interestingly, we observed a pattern of high diversity and divergence (Fig. 4a), particularly for genes within the recombinant blocks (Fig. 3b-e). The 38 (= top 1%) most divergent genes between *C. h. hominis* and *C. h. aquapotentis* were enriched in recombinant regions (χ2= 287.08, df = 1, p-value = 0.00049), with 37% present in these regions (Supplementary Table 5), compared to 0.9% of the other 1,645 divergent genes. These results suggest the genes undergoing frequent recombination accumulated beneficial polymorphisms that were maintained by balancing selection and that this increased population diversity.

Notably, the introgressed regions between *C. h. hominis* and *C. h. aquapotentis* included CHUDEA2_430 (*muc5*) for event 2 (chromosome 2), CHUDEA6_1080 (*gp60*) for event 3 (chromosome 6) and CHUDEA6_5270 (a hypothetical protein) for event 4 (chromosome 6). The introgression at CHUDEA6_5270 is particularly intriguing as it sheds further light on the evolution of *C. h. aquapotentis* and *C. h. hominis*, revealing how recent recombination events could be driving the virulence evolution of *C. h. hominis*. We identified two major CHUDEA6_5270 (hypothetical gene) haplotypes; Hap1 representing most *C. h. hominis* isolates, and Hap2 representing all *C. h. aquapotentis* plus a small subset of *C. h. hominis* (Fig. 5a), as well as many haplotypes associated only with *C. h. hominis*. No mutations were observed in Hap2, which is consistent with *C. h. aquapotentis* having evolved recently, being an estimated 392 years (29 – 1699 years; 5-95% CI) old, assuming a mutation rate of u=10^−8^ and 48h cell division time. Hap1 and Hap2 are diverged by 22 SNPs (Fig. 5b), which is unlikely to represent standing genetic variation, given this is 3-fold higher than the mean deviation among all other CHUDEA6_5270 haplotypes identified here. Instead, we propose CHUDEA6_5270-Hap2 might be an introgressed variant from a highly diverged, unsampled (sub)species that diverged from *C. hominis* around 2.36 million generations ago (assuming a mutation rate of u=10^−8^), which is equal to 12,742 (8,901 – 17,497 years; 5-95% CI) years (assuming 48h/replication). The emergence of CHUDEA6_5270-Hap2 in some *C. h. hominis* isolates (i.e., the red section of Hap2 in Fig. 6a) strongly implies that *C. h. aquapotentis* has introgressed (circa 400 years ago) into *C. h. hominis* leading to evolution under balancing selection (Fig. 5d).

**Fig 5.**
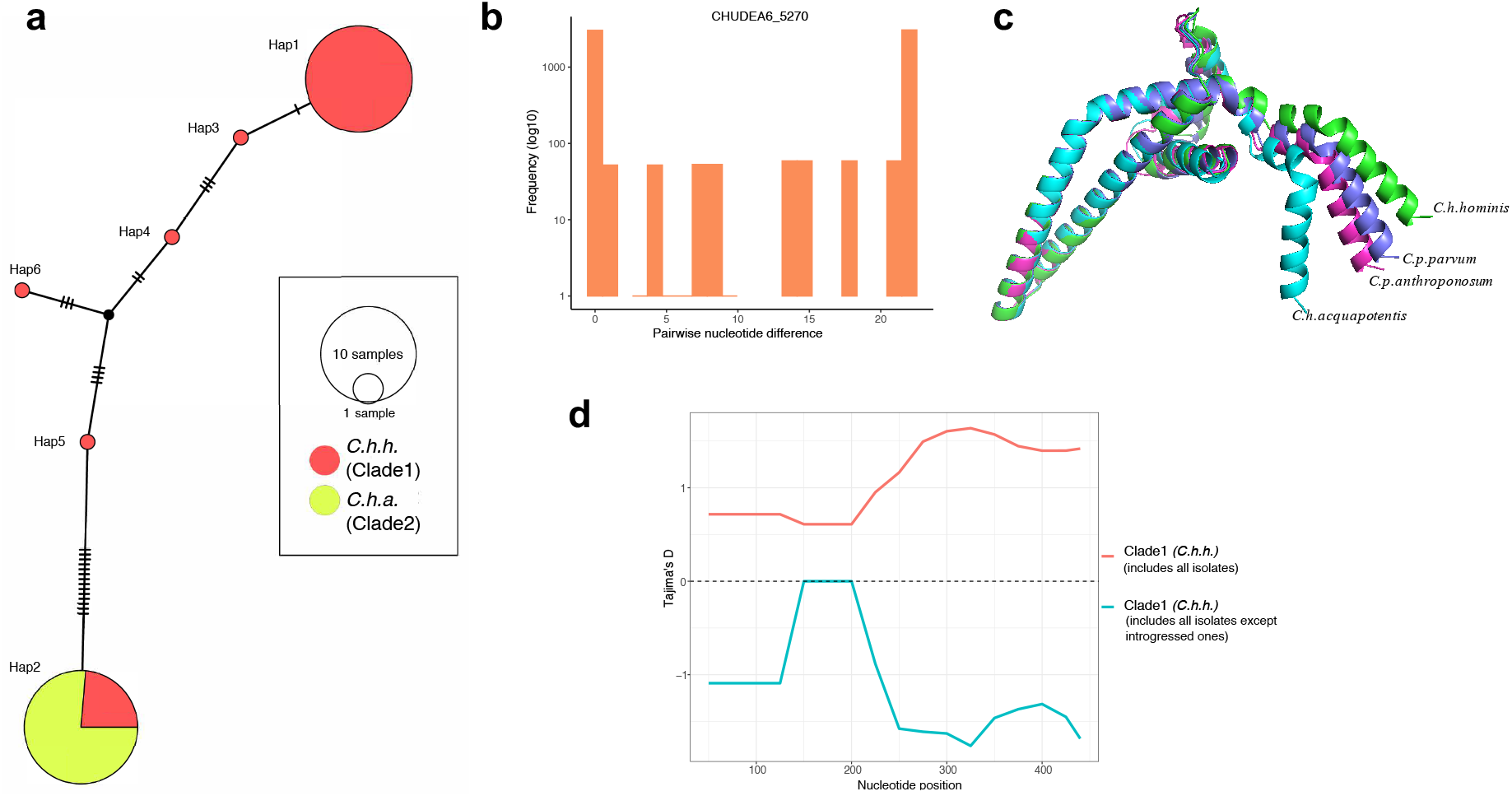
Illustrating diversifying selection between *C. hominis* subspecies and host adaptation at CHUDEA6_5270 (hypothetical gene). **a**. Haplotype network analyses illustrating haplotype diversification between *C. h. hominis* (clade 1) and *C. h. aquapotentis* (clade 2). **b**. Pairwise nucleotide divergence shows bimodal distribution, which, theoretically, can be explained both by balancing selection^79^, as well as by genetic introgression. **c**. Comparison of protein structure of CHUDEA6_5270 gene between *Cryptosporidium* species and subtypes demonstrates variation towards C.-terminal region. **d**. Introgressed-isolates driving balancing selection at gene CHUDEA6_5270 in *C. h. hominis*. Red line represents balancing selection (positive Tajima’s D) in *C. h. hominis* that also includes introgressed-isolates. Blue line represents purifying selection (negative Tajima’s D) in *C. h. hominis* after excluding introgressed-isolates.

We modelled the 3D protein structure of CHUDEA6_5270-Hap1 and Hap2, and compared these to similarly predicted 3D protein structures for the orthologous genes from *C. parvum parvum* and *C. parvum anthropanosum* (Fig. 5c). This modelling identified several conserved alpha-helices that appear to form a coiled-coil domain. CHUDEA6_5270 from *C*. *h. aquapotentis* encodes mutations near the C-terminal end, resulting in a notable kink that deviates from the other structures. This structural variation overlaps with an increase in Tajima’s D values toward the C-terminus of the protein, suggesting the region is under balancing selection (Fig. 5d).

A similar pattern is observed for CHUDEA2_430 (see Supplementary Fig. 6). However, given that both the subspecies are interspersed in the haplotype network, indicating against a specific directionality of introgression. This may indicate that CHUDEA_430 diverged with the divergence of the subspecies and continued to diversify. We were not able to generate a robust 3D structural model for CHUDEA2_430 or its *C. parvum* orthologs. We noted the gene encodes a large intrinsically disordered region (Supplementary Fig. 7) which is a consistent feature of mucin proteins^36^, whose structural confirmation is influenced by post-translation glycosylation^37^. Noting this, we did identify eight novel glycosylation sites in *C. h. aquapotentis* CHUDEA2_430 haplotypes not found in *C. h. hominis* (Supplementary Fig 5). Whether these impact interaction with host proteins is not known, but this would be consistent with glycosylated proteins in other pathogens^38^.

## Discussion

We examined the evolutionary genomics of a major human parasite, *C. hominis*, using 114 isolates from 16 countries across five continents. We identified two phylogenomic lineages within *C. hominis* that appear largely reproductively isolated, and we propose that these should be considered distinct subspecies. One *C. h. hominis* lineage comprises infections observed in low-to middle-income countries, mostly from Africa and Asia, which we propose be recognised as *C. hominis hominis*, referring to the fact that this subspecies represents the majority population. The second lineage (IbA10G2 subtype) includes isolates from high-income countries, namely Europe, North America and Oceania, which we suggest being named *C. hominis aquapotentis* (strong water), as this subspecies is predominant in countries with longstanding high sanitation and water quality indices.

These two subspecies are estimated to have been reproductively isolated for approximately 488 (84 – 2,199) years, except for the more recent genetic exchange at four genetic loci. The reproductive isolation coincides with improvements to sanitation in Europe. It is possible that *C. h. aquapotentis* evolved specialisations making it better suited to human infection allowing it to be more successful through direct transmission supported by a lower infectious dose. Such adaptations would have allowed *C. h. aquapotentis* to become dominant in higher-income countries where sanitation has reduced the level of environmental transmission. Indeed, epidemiological investigations in the UK identified direct person-to-person transmission as a key pathway for *C. h. aquapotentis*^24^. We hypothesise this may have resulted in its rapid population expansion over the last ∼500 years and (partial) reproductive isolation. In contrast, *C. h. hominis* experienced recent reduction in effective population size (*Ne*) within the past few decades. We detected two signatures of selective sweep in this subspecies, and we propose that this may have eroded some of the genetic variation, resulting in the marked drop in *Ne* = 5000 to *Ne* = 100 in the past.

Despite increased migration and international travelling in the past decades, our study suggests that there has been relatively little movement between continents for this parasite. This is in contrast to reports for *C. parvum*. Corsi *et. al*. recently found a higher proportion of admixture and gene flow between *C. parvum* populations and no evidence of population structuring by geographic region^39^. Despite the strong population sub-structuring in *C. hominis*, we found evidence of potential recombination and gene flow between the geographic populations and subspecies. We further investigated and identified the introgressed regions where we detected significant gene flow between the low- and high-income countries. Simulation-based analyses indicated this was most likely explained by ‘recent geneflow’ (circa 165 years ago). This would appear to be a secondary contact between the two subspecies after recent globalisation, illustrating higher migration rate from high-income to low-income countries, which facilitated gene flow, recombination, population admixture and selective sweep.

Finally, we have discovered genomic islands of putative virulence genes (GIPVs) contributing to population diversification between *C. h. aquapotentis* and *C. hominis hominis*. These islands have experienced relatively elevated recombination rate which has enriched nucleotide variation under balancing selection and the acquisition of non-synonymous SNPs, consistent with virulence factors driving host-parasite interactions. Intriguingly, the most significant signals within these analyses are driven by *gp60*, a hypothetical protein (CHUDEA6_5270) and a cluster of mucin-like genes (CHUDEA2_430, CHUDEA2_440 and CHUDEA2_450) found on chromosome 2. These genes are consistently identified as being under selection in the evolution of the *C. hominis* subspecies here and in similar observations made of *C. hominis* in Africa^40^. Their orthologs are associated with recombination between human-specific *C. parvum anthroponosum* relative to the zoonotic *C. parvum parvum* and appear to have driven convergence of the former with *C. hominis*^16^.

CHUDEA2_430 (*muc5*) and hypothetical protein CHUDEA6_5270 are the most notable, displaying significant diversifying selection between the two subspecies. Broadly, mucins mediate cell-cell interactions^41^, and modulate infectivity of *Cryptosporidium* sporozoites and merozoites and oocyst production^41,42^. In *C. parvum*, MUC5 is involved in host-cell invasion and an important determinant of host adaptation^41^ and highly expressed in the first 2 hours of infection in vitro^43^. MUC5 may also play a role in tethering the sporozoite to the oocyst wall^44^. Our analyses suggest *C. h. aquapotentis* and *C. h. hominis muc5* haplotypes diverged before or with the subspecies and subsequently diversified, which is consistent with prior observations implicating CHUDEA2_430 in the emergence of *C. h. aquapotentis*^10,45^. This appears to have resulted in the acquisition of novel glycosylation sites within *C. h. aquapotentis muc5* haplotypes. We cannot determine the functional consequence of these sites but note that glycosylation sites often mediate the specificity of mucin interactions with host proteins in a variety of pathogens^38^. In contrast, CHUDEA6_5270 displays a clear signal for the recent introgression of a novel *C. h. aquapotentis* haplotype into *C. h. hominis* after the divergence of these subspecies. This haplotype has notable, structurally relevant, mutations. Identifying the function of this gene, its potential role in infection and the relevance of the structural variation we have inferred here, should be considered a major research priority.

In conclusion, this work represents the first large scale population genomic study in any *Cryptosporidium* species, inferring the global population structure and evolutionary history of *C. hominis*. We propose recognition of two distinct subspecies, *C. h. hominis* and *C. h. aquapotentis*, with distinct demographic histories that have diverged circa 500 years ago. Although the subspecies differ in their global distribution, their gene pools are not completely isolated, and rare genetic exchanges have occurred in the recent past. We contend that many of the genes, CHUDEA2_430 and CHUDEA6_5270 in particular, in these introgression regions are involved in infection, and that their evolution in humans may be driving greater human specificity, virulence and transmissibility. It appears *C. h. aquapotentis* is playing a key role in this process, which is supported by previous observations based on multilocus typing^28^. This illustrates how human-mediated gene flow is involved in parasite evolution and genomic architecture, and how it could affect virulence evolution. Also, it shows that the GIPVs that result from population admixture in an anthroponotic species are under selection and involved in evolutionary arms race.

## Methods

### Parasite isolates

The *C. hominis* isolates newly sequenced for this study (n = 34) were archived stool samples collected at the *Cryptosporidium* Reference Unit in the UK. The species was determined by species-specific real-time PCR targeting the A135 gene^46^ and subtyped by PCR and sequencing of the *gp60* gene^47^. Supplementary Table 1 provides information about these isolates. Isolates were selected to mainly represent the dominant variant, IbA10G2, as defined by *gp60* sequencing.

### Processing of faecal samples for whole genome sequencing

Stool samples were processed as previously described^48^. Briefly, saturated salt-flotation was used to obtain a partially purified suspension of oocysts starting from 1-2 ml of each faecal sample. Oocysts were further purified from the suspension by immunomagnetic separation (IMS), using the Isolate® IMS kit (TCS Biosciences, Botolph Claydon, UK). IMS-purified oocysts were treated with bleach, and washed three times with nuclease-free water by centrifugation at 1,100X g for 5 min. The pellets were suspended in 200 μL of nuclease-free water for DNA extraction.

### DNA preparation and whole genome sequencing

Genomic DNA was extracted from purified *Cryptosporidium* oocysts by first performing eight cycles of freezing in liquid nitrogen for 1 min and thawing at 95°C for 1 min, and then using the QIAamp DNA extraction kit (Qiagen, Manchester, UK) according to the manufacturer’s instructions. The genomic DNA was eluted in 50 μL nuclease-free water, and the concentration measured using the Qubit dsDNA HS Assay Kit with the Qubit 1.0 fluorometer (Invitrogen, Paisley, UK), according to the manufacturer’s instructions.

Whole genome amplification (WGA) was performed using the Repli-g Midi kit (Qiagen, Milan, Italy), according to the manufacturer’s instructions. Briefly, 5 μL of genomic DNA (containing 1-10 ng of DNA) were mixed with 5 μL of denaturing solution and incubated at room temperature for 3 min. Next, 10 μL of stop solution were added to stabilise denatured DNA fragments. The reaction mixture was completed with 29 μL of buffer and 1 μL of phi29 polymerase, and allowed to proceed for 16 hours at 30°C. The reaction was stopped by heating at 63°C for 5 minutes. WGA products were visualised by electrophoresis on a 0.7% agarose gel, purified and quantified by Qubit as described above.

For Next Generation Sequencing (NGS) experiments, about 1 μg of purified WGA product was used to generate Illumina TruSeq 2x 150 bp paired-end libraries (average insert size: 500 bp), which were sequenced on an Illumina HiSeq 4000 platform (Illumina, SanDiego, CA). Library preparation and NGS experiments were performed by a commercial company (GATC, Germany).

### Whole genome global dataset

To perform a global comparative genomics of *C. hominis*, we supplemented our newly sequenced genome dataset by downloading all available published *C. hominis* genomes on till date (25^th^ July 2021), from the sequence read archive (SRA) of NCBI and from the EMBL’s European Nucleotide Archive (ENA) (see Supplementary Table 1). Collectively, these data represented 114 genomes from 16 countries across five continents.

### Data pre-processing and variant calling

Raw reads of the 114 *C. hominis* isolates were trimmed to remove adapter sequences and filtered for low-quality bases using Trimmomatic v.0.36^49^. The filtered reads were aligned to *C. hominis* UdeA01 reference genome^50,51^ using the maximal exact matches (MEM) algorithm implemented in Burrows-Wheeler Alignment (BWA) tool v.0.7^52^ with default settings. Sequence variants were called from the aligned reads of each isolate using the HaplotypeCaller method in the Genome Analysis Toolkit (GATK) v3.7.0^53^ as per GATK’s best practices pipeline^54^. Called variants were removed if quality depth (QD) < 2.0, Fisher strand (FS) > 60.0, mapping quality (MQ) < 40.0, mapping quality rank sum test (MQRankSum) < -12.5, read position rank sum test (ReadPosRankSum) < -8.0, Strand odds ratio (SOR) > 4.0, QUAL < 30, allele depth (AD) < 5, MAF < 0.05 or missingness > 0.5. Each of the 114 whole genomes assessed here had > 80% coverage of the *C. hominis* reference genome to at least the 5-fold depth. All identified variants were combined in one file and each isolate genotyped using the GenotypeGVCFs tool (GATK v3.7.0) ^53^ and the variants were separated into SNPs (i.e., nucleotide substitutions) and INDELs.

### Population genetic structure based on whole genome SNPs

The filtered SNPs were used for population structure, phylogenetic and clustering analyses. Samples having multiple co-infections were identified using estMOI^55^ and MOIMIX (https://github.com/bahlolab/moimix). The R package SNPRelate v.1.18^56^ was used for principal-component analysis (PCA) analysis. A maximum likelihood phylogenetic tree was constructed by IQ-TREE^57^ with 1000 bootstraps and visualised in iTOL v3^58^; the sister species *C. parvum* was used as an outgroup. We also constructed a consensus of 10^7^ trees using DensiTree 2^59^ in BEAST v2^60^. BEAST v2^60^ was also used to estimate the divergence time between the populations by using 95% highest posterior density (HPD); and SpeedDate (https://github.com/vanOosterhoutLab/SpeedDate.jl) to estimate the coalescence times between sequences by using 5-95% confidence interval (CI). We used mutation rate of 10^−8^ and a generation time of 48h/replication^16^ to date the coalescence times between sequences. A Neighbor-Net algorithm based network was generated using SplitsTree5^61^. Genetic structure was analysed by STRUCTURE v2.3 software^62^ for population number (K) ranging 2 - 10 and plotted by using plotSTR R package (https://github.com/DrewWham/Genetic-Structure-Tools). The optimal population genetic cluster value K was estimated by using CLUMPAK^63^.

### Population demographic history and divergence time estimation

We used Bayesian Markov Chain Monte Carlo (MCMC) method implemented in Beast v2 program^60^ to estimate the effective population size (*Ne*) of the *C. hominis* population. The nucleotide substitution model of HKY was selected. A strict molecular clock model and a Bayesian skyline coalescent tree prior was used with 10^9^ generations of MCMC chain and 10% burn-ins. Tracer v.1.7^64^ was used to assess chain convergence and effective sample size [ESS] > 200 and to construct the demographic history over time; i.e. Bayesian Skyline Plot (BSP). SweeD^30^ was used to detect windows of selective sweeps from genome-wide SNP dataset by using composite likelihood ratio (CLR) statistic that identifies signature of site frequency spectrum (SFS), with a grid size of 1000.

Demographic histories and migration rates between the *C. hominis* populations were estimated by using fastsimcoal2^65^ by using mutation rate of 10^−8^ and a generation time of 48h/replication^16^. We first inferred best parameters and the likelihoods for each of the demographic models – no geneflow, ongoing geneflow, early geneflow, recent geneflow and different geneflow, since the time of divergence (∼500 years) by running 100 independent iterations with 300,000 coalescent simulations and 60 optimisation cycles. Demographic model with the highest likelihood was then selected to run parameter estimation with block-bootstrapping of 100 replicates.

### Linkage, recombination and gene flow analyses

We inferred rate of decay of linkage disequilibrium (LD) by calculating the squared correlation of the coefficient (r^2^) between SNPs within 50kb using VCFtools^66^. LD blocks were also determined by calculating pairwise r^2^ between SNPs within chromosomes of each population. Recombination events were identified using the Recombination Detection Program version 4 (RDP4)^31^ using the RDP^67^, GENECONV^68^, BootScan^69^, MaxChi^70^ and Chimaera^71^ methods. Events were considered significant if at least three methods predicted their occurrence at a probability values, p ≤ 10^−5^. Recombination events with undetermined parental sequences were excluded from further HybridCheck analyses. Statistically significant recombination events were visualised and analysed using HybridCheck^33^ to determine the sequence similarity between the isolates involved in the events. HybridCheck program was also used to calculate the D statistic and estimate the gene flow between the populations.

### Population genetic and genomic analyses of coding region

Tajima’s D, Nucleotide diversity (π), Dxy and Fst were calculated using the PopGenome R-package^72^. Nonsynonymous (Ka) and synonymous (Ks) mutation rates were calculated by using Ka/Ks_Calculator^73^. Protein localisation (extracellular) was predicted using WoLF PSORT^74^ and information regarding predicted protein targeting (signalling peptides) genes were obtained from CryptoDB^50^. POPART program was used to generate haplotype networks^75^. AlphaFold was used to predict the protein structures^76^. Glycosylation sites were predicted by using NetNGlyc 4.0 Server^77^. Intrinsically disordered region in proteins were predicted using IUPred2A^78^. All statistical tests and results were performed and plotted in R (version 3.6.1).

## Data availability

All raw and processed sequencing data generated and analysed during the current study have been submitted to the NCBI Sequence Read Archive https://www.ncbi.nlm.nih.gov/bioproject/), under the BioProject PRJEB15112, PRJNA610731, PRJNA610732, PRJNA610735, PRJNA610737, PRJNA610738, PRJNA610739, PRJNA610740, PRJNA610741, PRJNA610742, PRJNA610743, PRJNA610744, PRJNA610745, PRJNA610746, PRJNA610747 and PRJNA610748.

## Acknowledgments

S.T. acknowledges the Australian Society for Parasitology (ASP) for a student conference travel grant, and JD Smyth Postgraduate Travel Award for a Network Researcher Exchange, Training and Travel Award; and Walter and Eliza Hall Research Institute (WEHI), Australia, for the top up scholarship. S.T. also acknowledges Namrata Srivastava (Monash University) for assisting with statistical data analysis.

## Author contributions

A.R.J., S.M.C., C.V.O. and S.T. conceived the study. A.R.J., S.M.C., C.V.O. and S.T. designed the analyses. S.M.C., R.M.C., G.R., D.E., and K.M.T. were involved in acquisition of data. S.T. performed the bioinformatics associated evolutionary genetic and genomic analyses. A.R.J., S.M.C., C.V.O. and S.T. wrote the manuscript. All authors read and approved the submission of the manuscript for the publication.

