## Supplementary information for "Global population genomics of two subspecies of *Cryptosporidium hominis* during 500 years of evolution"

**Title**

**Supplementary Text**

We undertook additional analyses of the highly admixed European isolate (UK_UKH4 of *gp60* subtype IaA14R3) that showed unique ancestry (Fig. 1B) and clustered with low-income countries (*C. h. hominis*). RDP4 identified four recombination events involving this admixed isolate in chromosomes 3, 4, 5 and 8 (Supplementary Fig. 3 and Supplementary Table 3). Moreover, a higher number of population clusters was determined in these chromosomes by the Evanno method (Supplementary Fig. 3), where the admixed isolate had shared ancestry with these clusters (Supplementary Fig. 4 & 5). The unique genomic evolution of this isolate is further evident in a SplitTree graph, which showed the presence of loops between *C. h. hominis* and *C. h. aquapotentis* (IbA10G2), and in a HybridCheck analysis, which revealed the signature of hybridisation and genetic introgression (Supplementary Fig. 4 & 5).

### Supplementary Table 1

List of the *C. hominis* isolates included in the study and available metadata.

| Sample code | Country origin | Run Accession | Bioproject | *gp60* subtype | Source | Collection year | Host | Recent Travel |
|---|---|---|---|---|---|---|---|---|
| Bangladesh_ERR2240054 | Bangladesh | ERR2240054 | PRJEB24168 | IdA15G1 | [1] | NA | Human | NA |
| Bangladesh_ERR2240055 | Bangladesh | ERR2240055 | PRJEB24168 | IdA15G1 | [1] | NA | Human | NA |
| Bangladesh_ERR2240056 | Bangladesh | ERR2240056 | PRJEB24168 | IaA25R3 | [1] | NA | Human | NA |
| Bangladesh_ERR2240057 | Bangladesh | ERR2240057 | PRJEB24168 | IaA25R3 | [1] | NA | Human | NA |
| Bangladesh_ERR2240058 | Bangladesh | ERR2240058 | PRJEB24168 | IaA18R3 | [1] | NA | Human | NA |
| Bangladesh_ERR2240059 | Bangladesh | ERR2240059 | PRJEB24168 | IbA9G3b | [1] | NA | Human | NA |
| Bangladesh_ERR2240060 | Bangladesh | ERR2240060 | PRJEB24168 | IbA9G3 | [1] | NA | Human | NA |
| Bangladesh_ERR2240061 | Bangladesh | ERR2240061 | PRJEB24168 | IaA18R3 | [1] | NA | Human | NA |
| Bangladesh_ERR2240062 | Bangladesh | ERR2240062 | PRJEB24168 | IaA18R3 | [1] | NA | Human | NA |
| Bangladesh_ERR2240063 | Bangladesh | ERR2240063 | PRJEB24168 | IfA16G1 | [1] | NA | Human | NA |
| Bangladesh_ERR2240064 | Bangladesh | ERR2240064 | PRJEB24168 | IbA9G3b | [1] | NA | Human | NA |
| Bangladesh_ERR2240065 | Bangladesh | ERR2240065 | PRJEB24168 | IfA9G3 | [1] | NA | Human | NA |
| Bangladesh_ERR2240066 | Bangladesh | ERR2240066 | PRJEB24168 | IbA9G3b | [1] | NA | Human | NA |
| Bangladesh_ERR2240067 | Bangladesh | ERR2240067 | PRJEB24168 | IaA18R3 | [1] | NA | Human | NA |
| Bangladesh_ERR2240068 | Bangladesh | ERR2240068 | PRJEB24168 | IbA9G3 | [1] | NA | Human | NA |
| Bangladesh_ERR2240069 | Bangladesh | ERR2240069 | PRJEB24168 | IfA13G1 | [1] | NA | Human | NA |
| Bangladesh_ERR2240070 | Bangladesh | ERR2240070 | PRJEB24168 | IaA18R3 | [1] | NA | Human | NA |
| Bangladesh_ERR2240071 | Bangladesh | ERR2240071 | PRJEB24168 | IaA26R3 | [1] | NA | Human | NA |
| Bangladesh_ERR2240072 | Bangladesh | ERR2240072 | PRJEB24168 | IaA25R3 | [1] | NA | Human | NA |
| Bangladesh_ERR2240073 | Bangladesh | ERR2240073 | PRJEB24168 | IeA11G3T3 | [1] | NA | Human | NA |
| Bangladesh_ERR2240074 | Bangladesh | ERR2240074 | PRJEB24168 | IeA11G3T3 | [1] | NA | Human | NA |
| Bangladesh_ERR2240075 | Bangladesh | ERR2240075 | PRJEB24168 | IaA19R3 | [1] | NA | Human | NA |
| Bangladesh_ERR2240076 | Bangladesh | ERR2240076 | PRJEB24168 | IaA18R3 | [1] | NA | Human | NA |
| Bangladesh_ERR2240077 | Bangladesh | ERR2240077 | PRJEB24168 | IbA9G3 | [1] | NA | Human | NA |
| Bangladesh_ERR2240078 | Bangladesh | ERR2240078 | PRJEB24168 | IbA9G3 | [1] | NA | Human | NA |
| Bangladesh_ERR2240079 | Bangladesh | ERR2240079 | PRJEB24168 | IaA27R3 | [1] | NA | Human | NA |
| Bangladesh_ERR2240080 | Bangladesh | ERR2240080 | PRJEB24168 | IaA27G3 | [1] | NA | Human | NA |
| Bangladesh_ERR2240081 | Bangladesh | ERR2240081 | PRJEB24168 | IdA14 | [1] | NA | Human | NA |
| Bangladesh_ERR2240082 | Bangladesh | ERR2240082 | PRJEB24168 | IaA19R3 | [1] | NA | Human | NA |
| Bangladesh_ERR2240083 | Bangladesh | ERR2240083 | PRJEB24168 | IbA9G3b | [1] | NA | Human | NA |
| Bangladesh_ERR2240084 | Bangladesh | ERR2240084 | PRJEB24168 | IaA27R3 | [1] | NA | Human | NA |
| Bangladesh_ERR2722072 | Bangladesh | ERR2722072 | PRJEB24168 | IaA18R3 | [1] | NA | Human | NA |
| India_SRR3098110 | Sweden | SRR3098110 | PRJNA307563 | IaA23R3 | [2] | 2013 | Human | India |
| Gabon_Afr30 | Gabon | SRR11843796 | PRJNA634705 | IdA22 | [3] | 2017 | Human | NA |
| Gabon_Afr33 | Gabon | SRR11843795 | PRJNA634705 | IbA9G3 | [3] | 2017 | Human | NA |
| Gabon_Afr34 | Gabon | SRR11843794 | PRJNA634705 | IbA13G3 | [3] | 2018 | Human | NA |

| | | | | | | | | |
|---|---|---|---|---|---|---|---|---|
| Gabon_Afr35 | Gabon | SRR11843793 | PRJNA634705 | IeA11G3T3 | [3] | 2017 | Human | NA |
| Ghana_Afr1 | Ghana | SRR11843813 | PRJNA634705 | IeA11G3T3 | [3] | 2016 | Human | NA |
| Ghana_Afr2 | Ghana | SRR11843812 | PRJNA634705 | IeA11G3T3 | [3] | 2016 | Human | NA |
| Ghana_Afr3 | Ghana | SRR11843801 | PRJNA634705 | IeA11G3T3 | [3] | 2016 | Human | NA |
| Ghana_Afr5 | Ghana | SRR11843792 | PRJNA634705 | IeA11G3T3 | [3] | 2016 | Human | NA |
| Ghana_Afr6 | Ghana | SRR11843791 | PRJNA634705 | IeA11G3T3 | [3] | 2016 | Human | NA |
| Ghana_Afr7 | Ghana | SRR11843790 | PRJNA634705 | IeA11G3T3 | [3] | 2016 | Human | NA |
| Ghana_Afr9 | Ghana | SRR11843789 | PRJNA634705 | IdA15 | [3] | 2016 | Human | NA |
| Madagascar_Afr11 | Madagascar | SRR11843787 | PRJNA634705 | IaA14 | [3] | 2017 | Human | NA |
| Madagascar_Afr12 | Madagascar | SRR11843786 | PRJNA634705 | IaA14 | [3] | 2017 | Human | NA |
| Madagascar_Afr13 | Madagascar | SRR11843811 | PRJNA634705 | IaA14 | [3] | 2017 | Human | NA |
| Madagascar_Afr10 | Madagascar | SRR11843788 | PRJNA634705 | IbA10G2 | [3] | 2017 | Human | NA |
| Ethiopia_SRR3098102 | Sweden | SRR3098102 | PRJNA307563 | IbA9G3 | [2] | 2014 | Human | Ethiopia |
| Kenya_SRR3098103 | Sweden | SRR3098103 | PRJNA307563 | IdA14 | [2] | 2012 | Human | Kenya |
| BurkinaF_SRR3098108 | Sweden | SRR3098108 | PRJNA307563 | IbA13G3 | [2] | 2013 | Human | Burkina Faso |
| Tanzania_SRR3098107 | Sweden | SRR3098107 | PRJNA307563 | IaA20R3 | [2] | 2013 | Human | Tanzania |
| Tanzania_Afr14 | Tanzania | SRR11843810 | PRJNA634705 | IfA14G1 | [3] | 2016 | Human | NA |
| Tanzania_Afr15 | Tanzania | SRR11843809 | PRJNA634705 | IdA17 | [3] | 2017 | Human | NA |
| Tanzania_Afr16 | Tanzania | SRR11843808 | PRJNA634705 | IdA17 | [3] | 2017 | Human | NA |
| Tanzania_Afr17 | Tanzania | SRR11843807 | PRJNA634705 | IeA11G3T3 | [3] | 2017 | Human | NA |
| Tanzania_Afr18 | Tanzania | SRR11843806 | PRJNA634705 | IdA17 | [3] | 2017 | Human | NA |
| Tanzania_Afr20 | Tanzania | SRR11843805 | PRJNA634705 | IaA19 | [3] | 2017 | Human | NA |
| Tanzania_Afr21 | Tanzania | SRR11843804 | PRJNA634705 | IeA11G3T3 | [3] | 2017 | Human | NA |
| Tanzania_Afr22 | Tanzania | SRR11843803 | PRJNA634705 | IeA11G3T3 | [3] | 2017 | Human | NA |
| Tanzania_Afr23 | Tanzania | SRR11843802 | PRJNA634705 | IaA25 | [3] | 2017 | Human | NA |
| Tanzania_Afr24 | Tanzania | SRR11843800 | PRJNA634705 | IdA17 | [3] | 2017 | Human | NA |
| Tanzania_Afr25 | Tanzania | SRR11843799 | PRJNA634705 | IeA11G3T3 | [3] | 2017 | Human | NA |
| Tanzania_Afr26 | Tanzania | SRR11843798 | PRJNA634705 | IbA9G3 | [3] | 2017 | Human | NA |
| UK_UKH4 | United Kingdom | SRR6143718 | PRJNA253838 | IaA14R3 | [4] | 2013 | Human | NA |
| UK_UKH6 | United Kingdom | SRR7895183 | PRJNA492838 | IdA30 | [5] | 2013 | Human | NA |
| UK_UKH30 | United Kingdom | SRR13222556 | PRJNA610741 | IbA9G3 | This study | 2015 | Human | Egypt |
| UK_UKH51 | United Kingdom | ERR2889329 | PRJEB15112 | IbA10G2 | This study | 2017 | Human | NA |
| UK_ UKH55 | United Kingdom | ERR2889330 | PRJEB15112 | IbA10G2 | This study | 2017 | Human | NA |
| UK_UKH56 | United Kingdom | ERR2889331 | PRJEB15112 | IbA10G2 | This study | 2017 | Human | NA |
| UK_UKH57 | United Kingdom | ERR2889332 | PRJEB15112 | IbA10G2 | This study | 2017 | Human | NA |
| UK_UKH58 | United Kingdom | ERR2889333 | PRJEB15112 | IbA10G2 | This study | 2017 | Human | NA |
| UK_UKH59 | United Kingdom | ERR2889334 | PRJEB15112 | IbA10G2 | This study | 2017 | Human | NA |
| UK_UKH60 | United Kingdom | ERR2889335 | PRJEB15112 | IbA10G2 | This study | 2017 | Human | NA |
| UK_UKH61 | United Kingdom | ERR2889336 | PRJEB15112 | IbA10G2 | This study | 2017 | Human | NA |
| UK_UKH62 | United Kingdom | ERR2889337 | PRJEB15112 | IbA10G2 | This study | 2017 | Human | NA |
| UK_UKH63 | United Kingdom | ERR2889338 | PRJEB15112 | IbA10G2 | This study | 2017 | Human | NA |
| UK_UKH64 | United Kingdom | ERR2889339 | PRJEB15112 | IbA10G2 | This study | 2017 | Human | NA |
| UK_UKH65 | United Kingdom | ERR2889340 | PRJEB15112 | IbA10G2 | This study | 2017 | Human | NA |

| | | | | | | | | |
|---|---|---|---|---|---|---|---|---|
| UK_UKH66 | United Kingdom | ERR2889341 | PRJEB15112 | IbA10G2 | This study | 2017 | Human | NA |
| UK_UKH67 | United Kingdom | ERR2889348 | PRJEB15112 | IbA10G2 | This study | 2017 | Human | NA |
| UK_UKH68 | United Kingdom | ERR2889349 | PRJEB15112 | IbA10G2 | This study | 2017 | Human | NA |
| UK_UKH69 | United Kingdom | ERR2889350 | PRJEB15112 | IbA10G2 | This study | 2017 | Human | NA |
| UK_UKH70 | United Kingdom | ERR2889351 | PRJEB15112 | IbA10G2 | This study | 2017 | Human | NA |
| UK_UKH71 | United Kingdom | ERR2889352 | PRJEB15112 | IbA10G2 | This study | 2017 | Human | NA |
| UK_UKH72 | United Kingdom | ERR2889353 | PRJEB15112 | IbA10G2 | This study | 2017 | Human | NA |
| UK_UKH3 | United Kingdom | SRR6131684 | PRJNA253834 | IbA10G2 | [4] | 2012 | Human | Turkey |
| UK_UKH5 | United Kingdom | SRR6144056 | PRJNA253839 | IbA10G2 | [4] | 2013 | Human | NA |
| UK_UKH23 | United Kingdom | SRR13222409 | PRJNA610731 | IbA10G2 | This study | 2015 | Human | Spain |
| UK_UKH24 | United Kingdom | SRR13222446 | PRJNA610732 | IbA10G2 | This study | 2015 | Human | Spain |
| UK_UKH25 | United Kingdom | SRR13222448 | PRJNA610735 | IbA10G2 | This study | 2015 | Human | Spain |
| UK_UKH26 | United Kingdom | SRR13222454 | PRJNA610737 | IbA10G2 | This study | 2015 | Human | Egypt |
| UK_UKH27 | United Kingdom | SRR13222458 | PRJNA610738 | IbA10G2 | This study | 2015 | Human | Egypt |
| UK_UKH28 | United Kingdom | SRR13222470 | PRJNA610739 | IbA10G2 | This study | 2015 | Human | Spain |
| UK_UKH29 | United Kingdom | SRR13222496 | PRJNA610740 | IbA10G2 | This study | 2015 | Human | Spain |
| UK_UKH31 | United Kingdom | SRR13222561 | PRJNA610742 | IbA10G2 | This study | 2015 | Human | Spain |
| UK_UKH32 | United Kingdom | SRR13222697 | PRJNA610743 | IbA10G2 | This study | 2015 | Human | NA |
| UK_UKH33 | United Kingdom | SRR13222828 | PRJNA610744 | IbA10G2 | This study | 2015 | Human | NA |
| UK_UKH34 | United Kingdom | SRR13223792 | PRJNA610745 | IbA10G2 | This study | 2015 | Human | NA |
| UK_UKH35 | United Kingdom | SRR13224181 | PRJNA610746 | IbA10G2 | This study | 2015 | Human | NA |
| UK_UKH36 | United Kingdom | SRR13224185 | PRJNA610747 | IbA10G2 | This study | 2015 | Human | NA |
| UK_UKH37 | United Kingdom | SRR13224632 | PRJNA610748 | IbA10G2 | This study | 2015 | Human | NA |
| Sweden_SRR3098109 | Sweden | SRR3098109 | PRJNA307563 | IfA12G1 | [2] | 2013 | Human | NA |
| Spain_SRR3098104 | Sweden | SRR3098104 | PRJNA307563 | IbA10G2 | [2] | 2013 | Human | Spain |
| Sweden_SRR3098106 | Sweden | SRR3098106 | PRJNA307563 | IbA10G2 | [2] | 2014 | Human | NA |
| Spain_SRR3098112 | Sweden | SRR3098112 | PRJNA307563 | IbA10G2 | [2] | 2014 | Human | Spain |
| Greece_SRR3098114 | Sweden | SRR3098114 | PRJNA307563 | IbA10G2 | [2] | 2014 | Human | Greece |
| USA_SRR1557959 | USA | SRR1557959 | PRJNA252787 | IaA28R4 | [6] | 2010 | Human | NA |
| USA_SRR1950894 | USA | SRR1950894 | PRJNA252787 | IaA28R4 | [6] | 2011 | Human | NA |
| USA_SRR1558150 | USA | SRR1558150 | PRJNA252787 | IbA10G2 | [6] | 2012 | Human | NA |
| USA_SRR1950893 | USA | SRR1950893 | PRJNA252787 | IbA10G2 | [6] | 2010 | Human | NA |
| Guatemala_SRR3098105 | Sweden | SRR3098105 | PRJNA307563 | IbA10G2 | [2] | 2014 | Human | Guatemala |
| NZ_SRR14089464 | New Zealand | SRR14089464 | PRJNA716090 | IbA10G2 | [7] | 2018 | Human | NA |
| NZ_SRR14089463 | New Zealand | SRR14089463 | PRJNA716090 | IbA10G2 | [7] | 2018 | Human | NA |

**Supplementary Table 2**

List of significant recombination events detected by RDP4[8] program between clade 1 and clade 2 in chromosomes 1, 2 and 6.

| Chrom | Event Number | Breakpoint Positions | | Recombinant Sequence(s) | Minor Parental Sequence(s) | Major Parental Sequence(s) | Detection methods | | | | |
|---|---|---|---|---|---|---|---|---|---|---|---|
| | | Begin | End | | | | RDP | GENECONV | Bootscan | Maxchi | Chimaera |
| 1 | 1 | 848362 | 855391 | Clade1 (n=4) | Unknown | Clade1 (n=65) | 1.72E-19 | 3.24E-17 | 3.42E-14 | 1.26E-04 | 2.48E-04 |
| 2 | 2 | 1346 | 11297 | Clade1 (n=17) | Clade2 (n=41) | Clade1 (n=55) | 3.77E-70 | 1.13E-64 | 3.83E-70 | 3.81E-23 | 3.36E-23 |
| | 2 | 1661 | 5361 | Clade2 (n=7) | Clade1 (n=50) | Clade2 (n=40) | 5.46E-27 | 2.02E-25 | 5.51E-27 | 1.35E-07 | 1.35E-07 |
| 6 | 3 | 261503 | 282666 | Clade1 (n=14) | Clade2 (n=45) | Clade1 (n=24) | 1.61E-71 | 2.93E-68 | 1.60E-71 | 2.56E-20 | 2.44E-20 |
| | 3 | 261249 | 263591 | Clade1 (n=13) | Clade2 (n=45) | Clade1 (n=22) | 4.32E-35 | 1.88E-32 | 2.60E-29 | 8.48E-10 | 1.29E-10 |
| | 3 | 259505 | 263543 | Clade1 (n=22) | Unknown | Clade1 (n=8) | 2.24E-25 | 3.87E-21 | 5.41E-26 | 2.84E-11 | 1.17E-10 |
| | 4 | 1209943 | 1300407 | Clade1 (n=10) | Clade2 (n=45) | Clade2 (n=46) | 7.98E-58 | 9.52E-56 | 9.53E-49 | 2.20E-17 | 1.45E-17 |
| | 4 | 1238761 | 1253435 | Clade1 (n=15) | Clade2 (n=45) | Clade1 (n=42) | 3.00E-43 | 6.84E-41 | 2.97E-43 | 2.17E-11 | 1.37E-11 |

### Supplementary Table 3

List of significant recombination events detected by RDP4[8] program in a highly admixed European isolate (UK_UKH4 of *gp60* subtype IaA14R3), in chromosomes 3, 4, 5 and 8.

| Chrom | Event Number | Breakpoint Positions | | Recombinant Sequence(s) | Minor Parental Sequence(s) | Major Parental Sequence(s) | Detection methods | | | | |
|---|---|---|---|---|---|---|---|---|---|---|---|
| | | Begin | End | | | | RDP | GENECONV | Bootscan | Maxchi | Chimaera |
| 3 | 1 | 478532 | 821544 | UK_UKH4 (Clade1) | Clade2 (n=45) | Clade1 (n=66) | 1.55E-47 | 4.75E-44 | 1.23E-47 | 1.43E-19 | 1.20E-19 |
| 4 | 2 | 97019 | 162083 | UK_UKH4 (Clade1) | Unknown | Clade1 (n=40) | 2.98E-07 | 3.29E-12 | 9.34E-14 | 4.57E-02 | 3.88E-02 |
| 5 | 3 | 99148 | 428100 | UK_UKH4 (Clade1) | Unknown | Clade1 (n=63) | 3.45E-28 | 8.67E-29 | 2.35E-19 | 1.85E-14 | 1.25E-14 |
| 8 | 1 | 666797 | 990157 | UK_UKH4 (Clade1) | Clade2 (n=45) | Clade1 (n=57) | 2.00E-54 | 6.80E-50 | 1.98E-54 | 2.92E-19 | 1.60E-18 |

### Supplementary Table 4

List of top 24 potential virulence genes and their associated population genetic metrices.

| Chrom | Chrom ID | Gene | Tajima's D | Nucleotide diversity | Dxy | Ka/Ks ratio | Recombination | Extracellular | Signal peptide |
|---|---|---|---|---|---|---|---|---|---|
| Chr2 | LN877948 | CHUDEA2_430 | 5.04924317 | 0.027263215 | 0.038396124 | 1.15774 | Yes | Yes | Yes |
| Chr6 | LN877952 | CHUDEA6_5270 | 4.83387277 | 0.02308583 | 0.036594203 | 0.637132 | Yes | Yes | Yes |
| Chr2 | LN877948 | CHUDEA2_440 | 4.74011177 | 0.02382591 | 0.034731335 | 0.836483 | Yes | Yes | Yes |
| Chr2 | LN877948 | CHUDEA2_450 | 4.56114583 | 0.013551459 | 0.019797529 | 0.461748 | Yes | Yes | No |
| Chr6 | LN877952 | CHUDEA6_1080 | 2.67308457 | 0.008568619 | 0.009621962 | 0.759268 | Yes | Yes | Yes |
| Chr6 | LN877952 | CHUDEA6_1030 | 4.05646271 | 0.003911115 | 0.007853953 | 3.17027 | Yes | Yes | Yes |
| Chr6 | LN877952 | CHUDEA6_5260 | 3.8875367 | 0.002501751 | 0.003890499 | 0.241018 | Yes | No | No |
| Chr6 | LN877952 | CHUDEA6_1100 | 3.36852674 | 0.001768587 | 0.003322986 | 0.349574 | Yes | Yes | Yes |
| Chr7 | LN877953 | CHUDEA7_200 | 2.69090993 | 0.001356426 | 0.00274301 | 0.28405 | No | Yes | Yes |
| Chr2 | LN877948 | CHUDEA2_3270 | 3.41620427 | 0.001453499 | 0.002703619 | 0.193419 | No | No | No |
| Chr7 | LN877953 | CHUDEA7_650 | 2.79403816 | 0.001281769 | 0.002460284 | 0.290335 | No | No | No |
| Chr6 | LN877952 | CHUDEA6_2830 | 2.50950496 | 0.001553752 | 0.002437824 | 0.276323 | No | No | No |
| Chr6 | LN877952 | CHUDEA6_4520 | 2.75593862 | 0.000957408 | 0.00195719 | 0.568518 | No | No | No |
| Chr7 | LN877953 | CHUDEA7_580 | 2.91518247 | 0.000945169 | 0.001809445 | 0.557577 | No | No | No |
| Chr7 | LN877953 | CHUDEA7_1380 | 2.50146676 | 0.00099094 | 0.001715119 | 0.285047 | No | No | Yes |
| Chr5 | LN877951 | CHUDEA5_1120 | 2.79736431 | 0.000848783 | 0.001685789 | 0.524253 | No | No | No |
| Chr6 | LN877952 | CHUDEA6_4910 | 2.80314464 | 0.000833936 | 0.001604819 | 0.473719 | No | No | No |
| Chr3 | LN877949 | CHUDEA3_4260 | 2.49564189 | 0.001069508 | 0.001567676 | 0.527893 | No | No | No |
| Chr8 | LN877954 | CHUDEA8_810 | 2.73791027 | 0.000805645 | 0.001549064 | 1.34268 | No | No | No |
| Chr5 | LN877951 | CHUDEA5_4040 | 2.90614675 | 0.000755103 | 0.00150644 | 0.734507 | No | No | No |
| Chr1 | LN877947 | CHUDEA1_3450 | 2.63999445 | 0.000897162 | 0.001482651 | 0.367788 | No | No | No |
| Chr5 | LN877951 | CHUDEA5_2000 | 2.76701759 | 0.000727004 | 0.00147531 | 0.5015 | No | No | No |
| Chr1 | LN877947 | CHUDEA1_3540 | 3.30908675 | 0.000732585 | 0.001374333 | 1.02037 | No | No | No |
| Chr1 | LN877947 | CHUDEA1_260 | 2.51197826 | 0.000730038 | 0.001228733 | 0.244469 | No | Yes | Yes |

**Supplementary Table 5**

List of 38 most divergent genes (top 1% genes) and their associated population genetic metrices. These genes were significantly enriched in recombinant regions ($\chi2$= 287.08, df = 1, p-value = 0.00049) between *C. h. hominis* and *C. h. aquapotentis*.

| Chrom | Chrom ID | Gene | Rank | Tajima's D | Nucleotide diversity | Dxy | Recombination |
|---|---|---|---|---|---|---|---|
| Chr2 | LN877948 | CHUDEA2_430 | 1 | 5.04924317 | 0.02726322 | 0.03839612 | Yes |
| Chr6 | LN877952 | CHUDEA6_5270 | 2 | 4.83387277 | 0.02308583 | 0.0365942 | Yes |
| Chr2 | LN877948 | CHUDEA2_440 | 3 | 4.74011177 | 0.02382591 | 0.03473134 | Yes |
| Chr2 | LN877948 | CHUDEA2_450 | 4 | 4.56114583 | 0.01355146 | 0.01979753 | Yes |
| Chr6 | LN877952 | CHUDEA6_1000 | 5 | 2.19231838 | 0.00611982 | 0.00982528 | Yes |
| Chr6 | LN877952 | CHUDEA6_1080 | 6 | 2.67308457 | 0.00856862 | 0.00962196 | Yes |
| Chr1 | LN877947 | CHUDEA1_3810 | 7 | 1.81524237 | 0.00439554 | 0.00808404 | Yes |
| Chr6 | LN877952 | CHUDEA6_1030 | 8 | 4.05646271 | 0.00391112 | 0.00785395 | Yes |
| Chr8 | LN877954 | CHUDEA8_2140 | 9 | 1.74227725 | 0.00286945 | 0.00595238 | No |
| Chr1 | LN877947 | CHUDEA1_900 | 10 | 1.80029845 | 0.0032147 | 0.00560765 | No |
| Chr6 | LN877952 | CHUDEA6_1070 | 11 | 2.60189871 | 0.00409269 | 0.00491202 | Yes |
| Chr5 | LN877951 | CHUDEA5_new_05 | 12 | 1.6092262 | 0.0025025 | 0.00404125 | No |
| Chr6 | LN877952 | CHUDEA6_1020 | 13 | 2.6446673 | 0.00196873 | 0.00395062 | Yes |
| Chr6 | LN877952 | CHUDEA6_5260 | 14 | 3.8875367 | 0.00250175 | 0.0038905 | Yes |
| Chr6 | LN877952 | CHUDEA6_4450 | 15 | 1.74227725 | 0.00169147 | 0.00350877 | No |
| Chr2 | LN877948 | CHUDEA2_620 | 16 | 1.7634596 | 0.00172212 | 0.00349471 | No |
| Chr4 | LN877950 | CHUDEA4_1220 | 17 | 2.17922328 | 0.00202796 | 0.00349002 | No |
| Chr4 | LN877950 | CHUDEA4_2500 | 18 | 2.42788628 | 0.00180648 | 0.0034772 | No |
| Chr6 | LN877952 | CHUDEA6_60 | 19 | 3.90999432 | 0.00172457 | 0.00339283 | No |
| Chr1 | LN877947 | CHUDEA1_1370 | 20 | 2.3567791 | 0.00179349 | 0.00339238 | No |
| Chr6 | LN877952 | CHUDEA6_1100 | 21 | 3.36852674 | 0.00176859 | 0.00332299 | Yes |
| Chr6 | LN877952 | CHUDEA6_150 | 22 | 1.95663044 | 0.00171608 | 0.00329839 | No |
| Chr1 | LN877947 | CHUDEA1_2690 | 23 | 1.66767857 | 0.00163018 | 0.00324074 | No |
| Chr6 | LN877952 | CHUDEA6_970 | 24 | 2.50738449 | 0.00162321 | 0.00312582 | Yes |
| Chr2 | LN877948 | CHUDEA2_390 | 25 | 1.99196834 | 0.00189801 | 0.00302403 | No |
| Chr7 | LN877953 | CHUDEA7_4040 | 26 | 1.80029845 | 0.0015467 | 0.00300793 | No |
| Chr7 | LN877953 | CHUDEA7_4430 | 27 | 1.7828 | 0.00150895 | 0.00299696 | No |
| Chr4 | LN877950 | CHUDEA4_1040 | 28 | 2.40562664 | 0.0015135 | 0.00297459 | No |
| Chr6 | LN877952 | CHUDEA6_1360 | 29 | 2.18055695 | 0.00151763 | 0.00295567 | No |
| Chr2 | LN877948 | CHUDEA2_1070 | 30 | 1.81595497 | 0.001555 | 0.00285703 | No |
| Chr5 | LN877951 | CHUDEA5_2580 | 31 | 2.36790892 | 0.00138953 | 0.00281976 | No |
| Chr8 | LN877954 | CHUDEA8_4700 | 32 | 1.7828 | 0.00139288 | 0.00276642 | No |

| | | | | | | | |
|---|---|---|---|---|---|---|---|
| Chr7 | LN877953 | CHUDEA7_200 | 33 | 2.69090993 | 0.00135643 | 0.00274301 | No |
| Chr2 | LN877948 | CHUDEA2_3270 | 34 | 3.41620427 | 0.0014535 | 0.00270362 | No |
| Chr5 | LN877951 | CHUDEA5_1970 | 35 | 1.80029845 | 0.00136625 | 0.00265701 | No |
| Chr6 | LN877952 | CHUDEA6_5280 | 36 | 3.67033242 | 0.00174592 | 0.00260826 | Yes |
| Chr6 | LN877952 | CHUDEA6_4640 | 37 | 1.80029845 | 0.0013116 | 0.00255073 | No |
| Chr4 | LN877950 | CHUDEA4_3220 | 38 | 1.74227725 | 0.00119029 | 0.00246914 | No |

**Supplementary Fig 1**

Identification of regions with selective sweep. Plot of composite likelihood ratio (CLR) across *C. hominis* genome where red and blue dotted lines are associated with *C. h. hominis* (clade1) and *C. h. acquapotentis* (clade2), respectively. Horizontal dashed black line represents cutoff of 99.99 percentile. Two genomic regions with highest peaks of CLR (> 99.99 percentile) are observed in chromosome 6 in *C. h. hominis*.

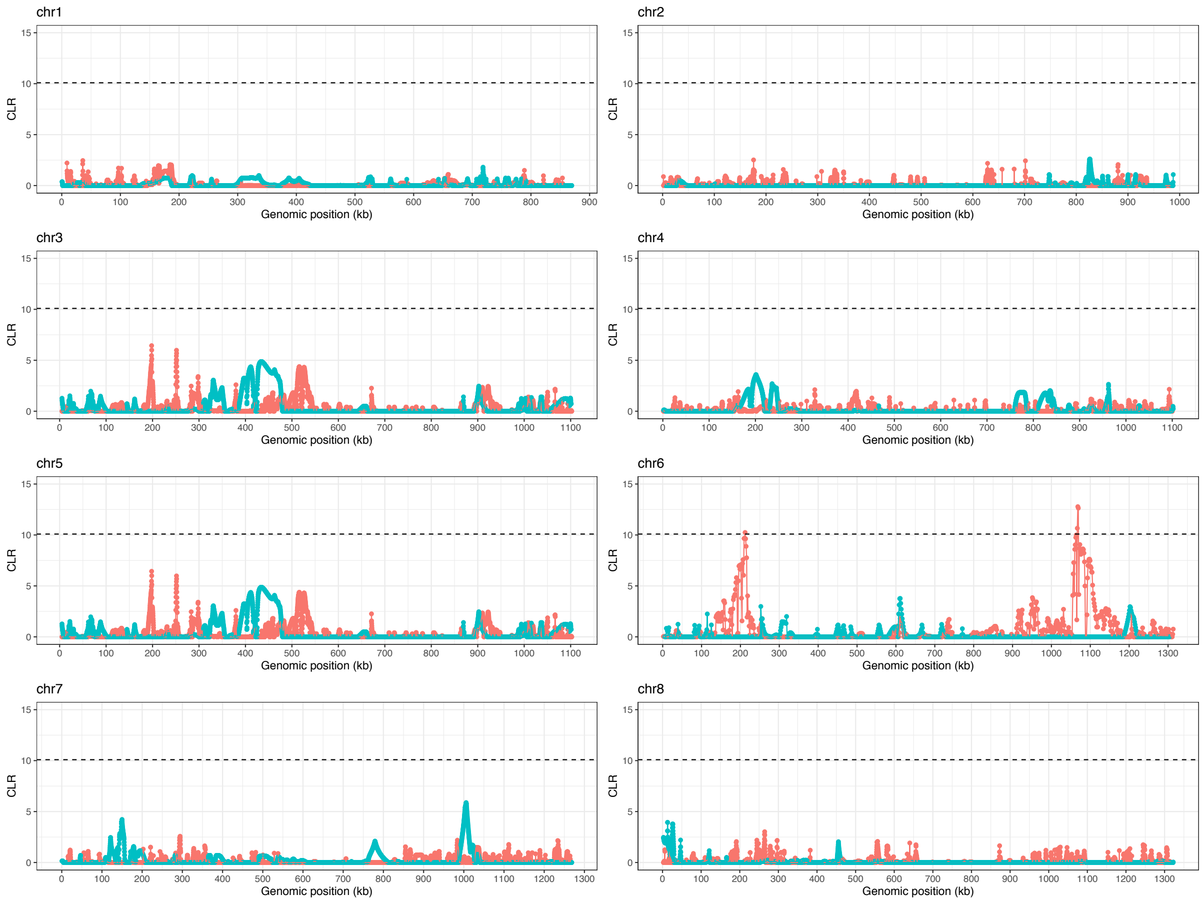

**Supplementary Fig 2**

Signature of selective sweep in chromosome 6 of *C. h. hominis*. Panels represent composite likelihood ratio (CLR) across chromosome 6 (top), nucleotide diversity (middle) and Tajima's D values (bottom). Windows of selective sweeps (CLR > 99.99 percentile) are highlighted with dashed boxes and genomic positions of recombinant events are illustrated with solid rectangular boxes.

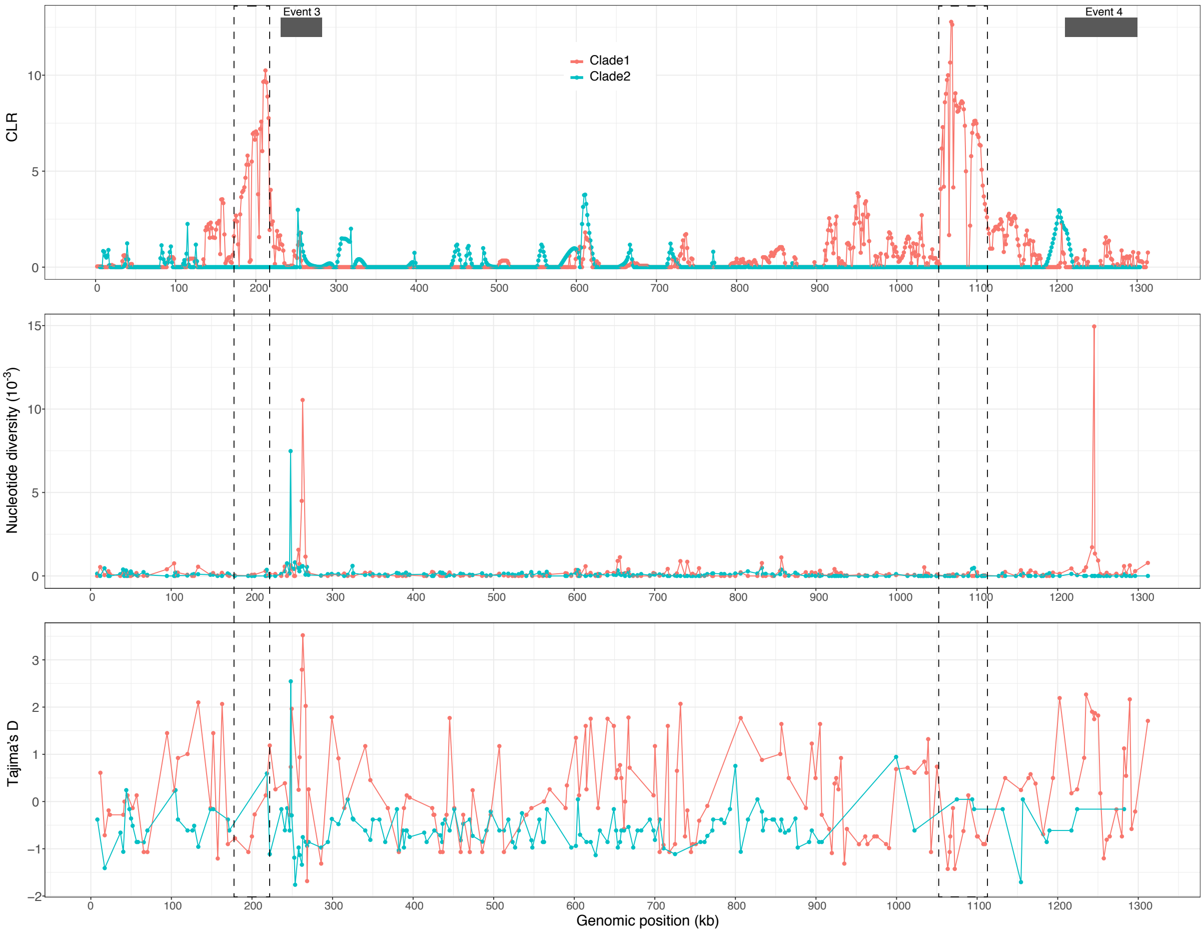

**Supplementary Fig 3**

Illustration of recombination and population structure in an admixed isolate UK_UKH4. **a.** The positions of four recombination events are indicated by black rectangles. **b**. Best K values determined by Evanno method[9] for each chromosome.

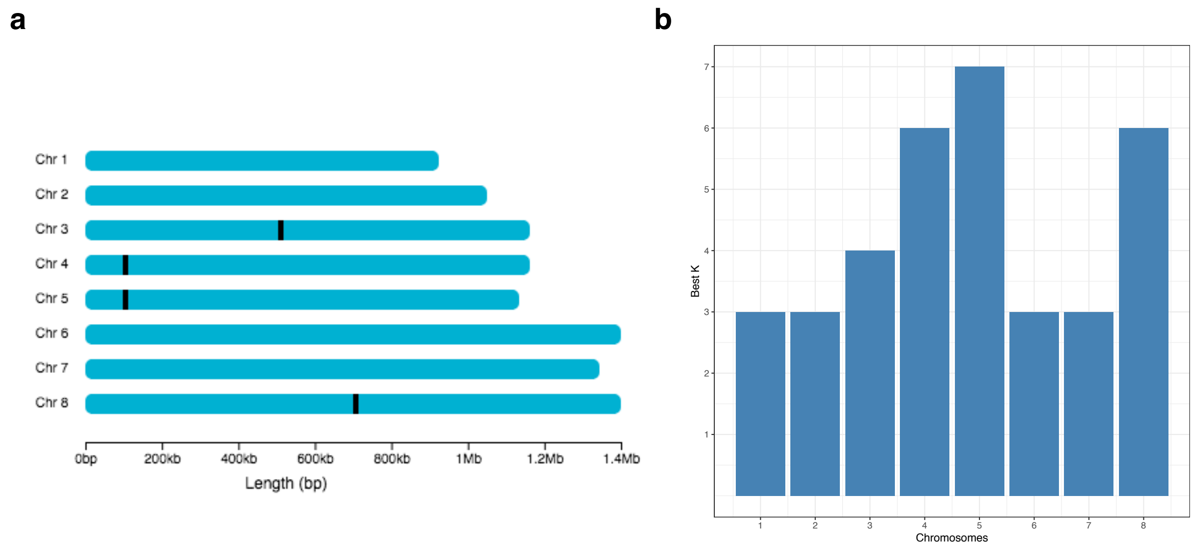

### Supplementary Fig 4

Admixed nature of an isolate UK_UKH4 revealed through population structure and introgression analyses in chromosome 3. **a**. Structure graph showing population clusters (UK_UKH4 is indicated by an arrow). **b.** Splitstree showing the position of the admixed isolate in between the two clades. **c**. HybridCheck graph illustrating the signature of introgression in UK_UKH4 where clade 1 (e.g., Tanzania_Afr14) and clade 2 (e.g., NZ_SRR14089463) isolates represents major and minor parental sequences, respectively, as detected by the RDP4 program.

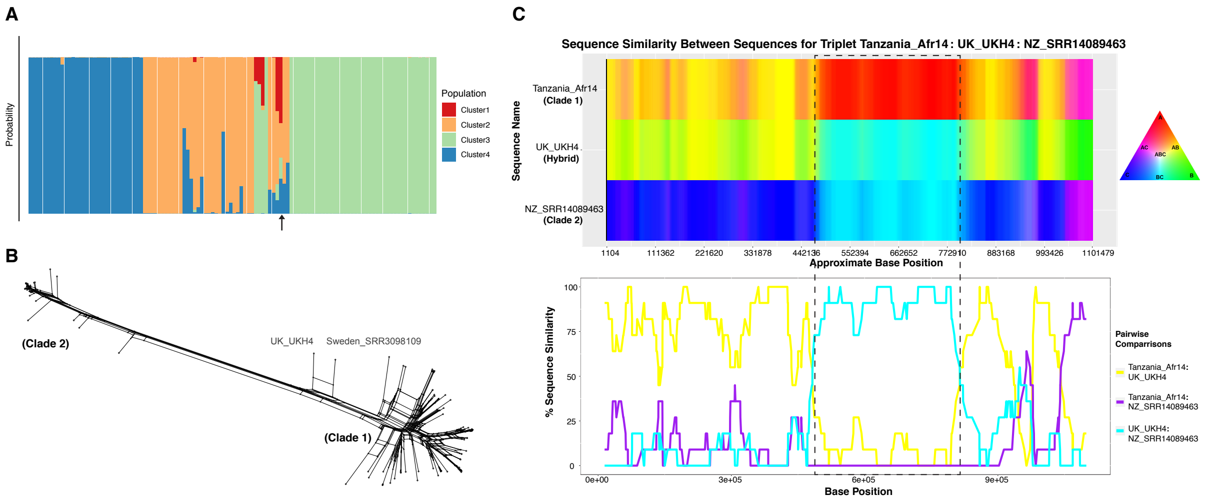

### Supplementary Fig 5

Admixed nature of an isolate UK_UKH4 revealed through population structure and introgression analyses in chromosome 8. **a**. Structure graph showing population clusters (UK_UKH4 is indicated by an arrow). **b.** Splitstree showing the position of the admixed isolate in between the two clades. **c**. HybridCheck graph illustrating the signature of introgression in UK_UKH4 where clade 1 (e.g., Bangladesh_ERR2240065) and clade 2 (e.g., UK_UKH3) isolates represents major and minor parental sequences, respectively, as detected by the RDP4 program.

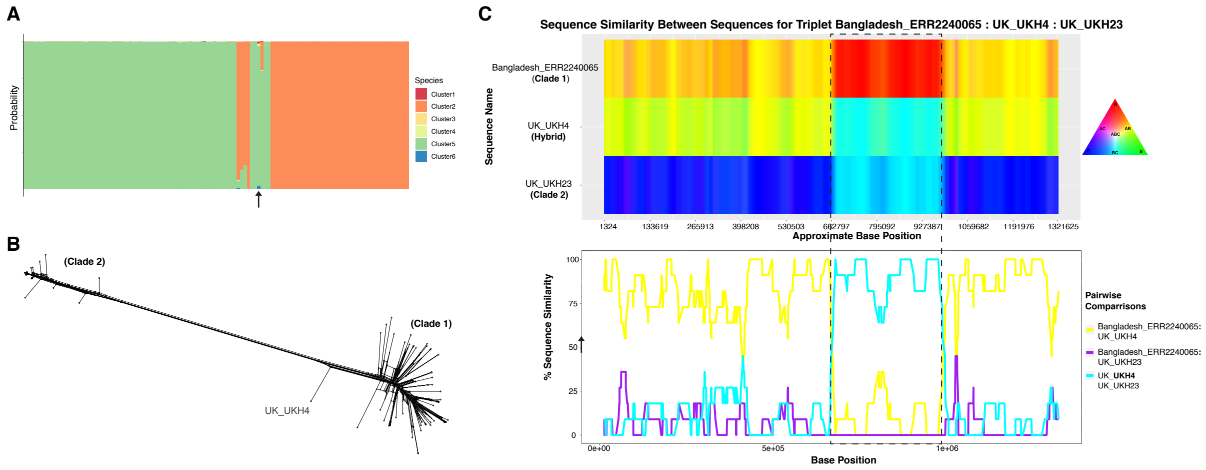

### Supplementary Fig 6

Illustrating diversifying selection between *C. hominis* subspecies and host adaptation at CHUDEA2_430 (MUC5). **a**. Haplotype network analyses of CHUDEA2_430 gene, illustrating haplotype diversification between cade 1 and clade 2. The divergence between *C.h.h.* (clade1) and *C.h.a.* (clade2) at the shared haplotype (Hap2) is estimated to be 316 [23 – 1,365 years; 5-95% CI] years. **b**. Pairwise nucleotide divergence shows bimodal distribution in CHUDEA2_430 gene, which, theoretically, can be explained both by balancing selection[10], as well as by genetic introgression. **c**. Introgressed-isolates driving balancing selection at gene CHUDEA2_430 in *C. h. hominis*. Red line represents balancing selection (positive Tajima's D) in *C. h. hominis* that also includes introgressed-isolates. Blue line represents purifying selection (negative Tajima's D) in *C. h. hominis* after excluding introgressed-isolates. **d**. Differences in glycosylation sites between *C. h. hominis* and *C. h. acquapotentis*.

a

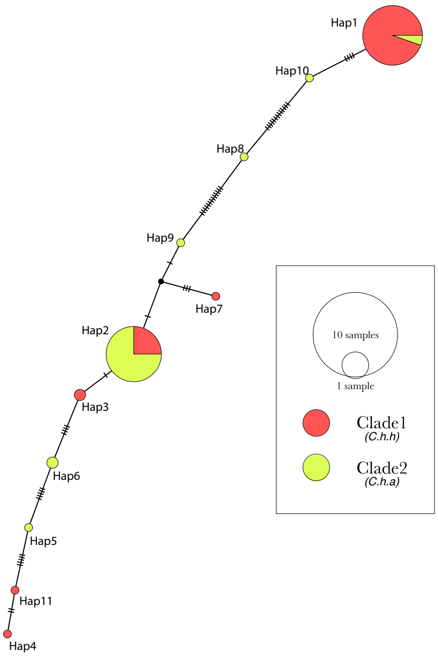


b

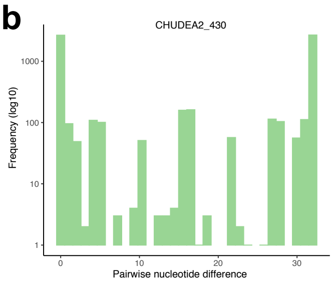


d

| Amino Acid | *C. h. acquapotentis* | *C. h. hominis* |
|---|---|---|
| 44 | Positive | - |
| 78 | - | Positive |
| 79 | Positive | - |
| 95 | Positive | - |
| 158 | Positive | - |
| 164 | Positive | - |
| 178 | - | Positive |
| 179 | Positive | - |
| 187 | Positive | - |
| 191 | Positive | - |

c

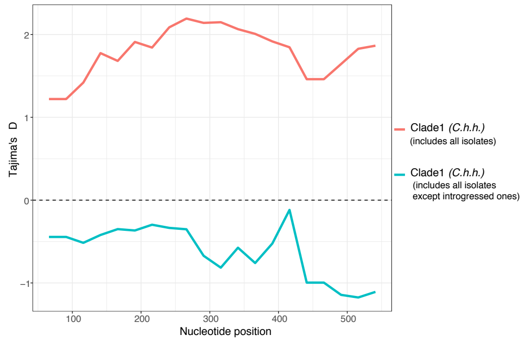

**Supplementary Fig 7**

Intrinsically disordered region in CHUDEA2_430 (MUC5) encoded protein using IUPred2A[11].

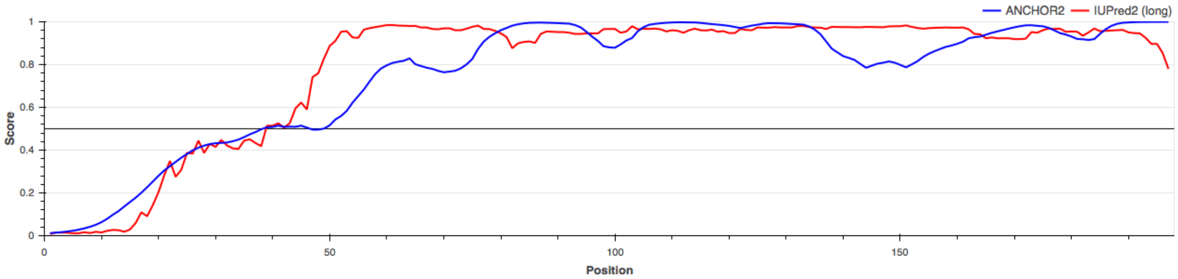
